# A viral histone-like protein exploits antagonism between linker histones and HMGB proteins to obstruct the cell cycle

**DOI:** 10.1101/2020.12.11.418129

**Authors:** Kelsey L. Lynch, Mongoljin Bat-Erdene, Melanie R. Dillon, Hannah C. Lewis, Daphne C. Avgousti

## Abstract

Virus infection necessarily requires redirecting cellular resources towards viral progeny production. Adenovirus encodes the histone-like protein VII that causes catastrophic global reorganization of host chromatin to promote virus infection. Protein VII recruits the family of high mobility group box (HMGB) proteins to chromatin along with the histone chaperone SET. As a consequence of this recruitment, we find that protein VII causes chromatin-depletion of several linker histone H1 isoforms. The relationship between linker histone H1 and the functionally opposite HMGB proteins is critical for higher order chromatin structure. However, the physiological consequences of perturbing this relationship are largely unknown. Here, we employ complementary systems in *Saccharomyces cerevisiae* and human cells to demonstrate that adenovirus protein VII disrupts the H1-HMGB balance to obstruct the cell cycle. We find that protein VII causes an accumulation of G2/M cells both in yeast and human systems, underscoring the high conservation of this chromatin vulnerability. In contrast, adenovirus E1A and E1B proteins are well-established to override cell cycle regulation and promote transformation of human cells. Strikingly, we find that protein VII obstructs the cell cycle even in the presence of E1A and E1B, suggesting that protein VII-directed cell cycle disruption ensures host resources are directed towards viral proliferation during infection. Together, our results demonstrate that protein VII targets H1-HMGB1 antagonism to obstruct cell cycle progression, revealing an unexpected chromatin vulnerability exploited for viral benefit.

## Introduction

Viral takeover of cellular processes is essential for the success of these intracellular pathogens. Controlling host chromatin is an integral aspect of viral infection that includes re-directing host resources for transcription, DNA replication, and cell division towards viral replication. Several examples have recently come to light that highlight viral alteration to host gene expression through modifications of core histone tails^1–7^. Host chromatin may also be manipulated through displacement of linker histones. Linker histones are critical for genome compaction^8–10^ and compete for binding sites with high mobility group (HMG) proteins^11^, which in contrast promote decompaction of chromatin and increase DNA accessibility^12–15^. This interplay between linker histones and HMG proteins is a vulnerability that may be a target of viral manipulation.

The adenovirus genome is packaged with a small basic protein known as protein VII^16–18^. This 20 kDa core protein forms a ‘beads on a string’ structure within virus particles and recruits the histone chaperone protein SET^19–21^ onto viral genomes as they enter the nucleus. SET is thought to populate the viral genomes with histones to promote viral gene expression^22–25^. The first viral gene products, E1A and E1B, are then expressed leading to subsequent viral gene activation, global changes in host histone modifications, and bypass of cell cycle checkpoints^26,28^. E1A has been the focus of many studies of cellular transformation and epigenetic reprogramming by viruses^29,30^, although it remains unclear why transformation does not occur in humans as a result of adenovirus infection^26,31^. Protein VII, along with other viral structural proteins, is transcribed late during infection^28^. In addition to its structural role inside virion cores, protein VII localizes to host chromatin and disrupts normal nuclear processes^32,33^. Protein VII is post-translationally modified and these modifications regulate protein VII’s localization to chromatin^32,33^. Recombinant protein VII binds purified nucleosomes directly, protecting approximately 165 bp of DNA from digestion by micrococcal nuclease, suggesting that it may have similar binding sites as the linker histones^32^.

Several host proteins are significantly enriched in chromatin in the presence of protein VII as measured by salt fractionation followed by mass spectrometry^32,34^ including SET and the family of high mobility group box (HMGB)^14^ proteins. Without protein VII, SET and HMGB proteins are transiently bound to chromatin. SET, also known as TAF-Iß, has been described as a linker histone chaperone^36,37^ while HMGB1 has been shown to act as an antagonist of histone H1^12,13,15^. Here, we use complementary systems in *Saccharomyces cerevisiae* and human cells to show that adenovirus protein VII requires HMGB1 and SET to disrupt chromatin and cause cell growth defects. We determine that these deficiencies in growth are due to deviations in cell cycle progression. We find that protein VII can slow cell cycle progression even in the presence of adenovirus E1A, which normally leads to a bypass of cell cycle checkpoints thereby promoting transformation. Therefore, we propose that during adenovirus infection protein VII obstructs mitosis to ensure cellular resources are directed towards viral progeny production rather than cell division. This study is the first to show viral exploitation of the H1-HMGB1 interaction and subsequent delay of the cell cycle for viral benefit.

## Results

### Human adenovirus protein VII causes growth defects in budding yeast

In addition to the enrichment of SET and HMGBs in chromatin upon protein VII expression in human lung epithelial cells, we discovered that several isoforms of the linker histone H1 were significantly depleted from chromatin in the presence of protein VII (Supplemental Figure 1a). Depletion of linker histones has recently come to light as a regulatory mechanism for diverse cellular outcomes^38–40^. Thus, our surprising finding led us to hypothesize that protein VII harnesses the antagonism between HMGB1 and H1 by co-opting SET and HMGB1 to displace histone H1 in host chromatin (Supplemental Figure 1b). We decided to test our model with genetic studies in *Saccharomyces cerevisiae,* which faithfully represents many aspects of mammalian chromatin with less genetic redundancy than mammalian cells^41,42^.

We generated a selectable, inducible expression plasmid bearing a yeast codon-optimized version of mature protein VII from adenovirus type 5 (Supplemental Figure 1c). We introduced the inducible expression plasmid into the wild type (WT) lab strain W303 and assayed growth on solid media using a serial dilution growth assay (Figure 1a). Strikingly, we found that expression of protein VII in budding yeast resulted in a ~40% growth deficit compared to cells expressing GFP (Figure 1b). Furthermore, upon mutation of five post-translationally modified (PTM) sites previously found to impact protein VII’s localization to chromatin^32,33^, the growth defect was rescued (Figure 1a and b). We further characterized the growth defect by inducing expression during log phase growth in liquid culture and calculating the exponential growth rate. Without the induction of expression, there was no difference in exponential growth rate among the three strains (Figure 1c). However, expression of protein VII caused a significant reduction in growth compared to GFP-expressing cells. The growth defect was rescued upon mutation of protein VII and the exponential growth rate of cells expressing VII-ΔPTM was comparable to GFP. Since protein expression is induced by galactose, a less-preferred energy source, the absolute growth rates for all induced strains were lower than the same strains grown in dextrose media without transgene expression, as expected. We confirmed comparable expression of each protein during exponential growth by western blot (Figure 1d). Furthermore, we found that the growth defect caused by protein VII was not unique to the W303 strain as we observed comparable results in an alternate lab strain, FY602 (Figure1e-h). The similar outcomes in both strains highlight the robust and conserved nature of protein VII’s impact on yeast.

**Figure 1.**
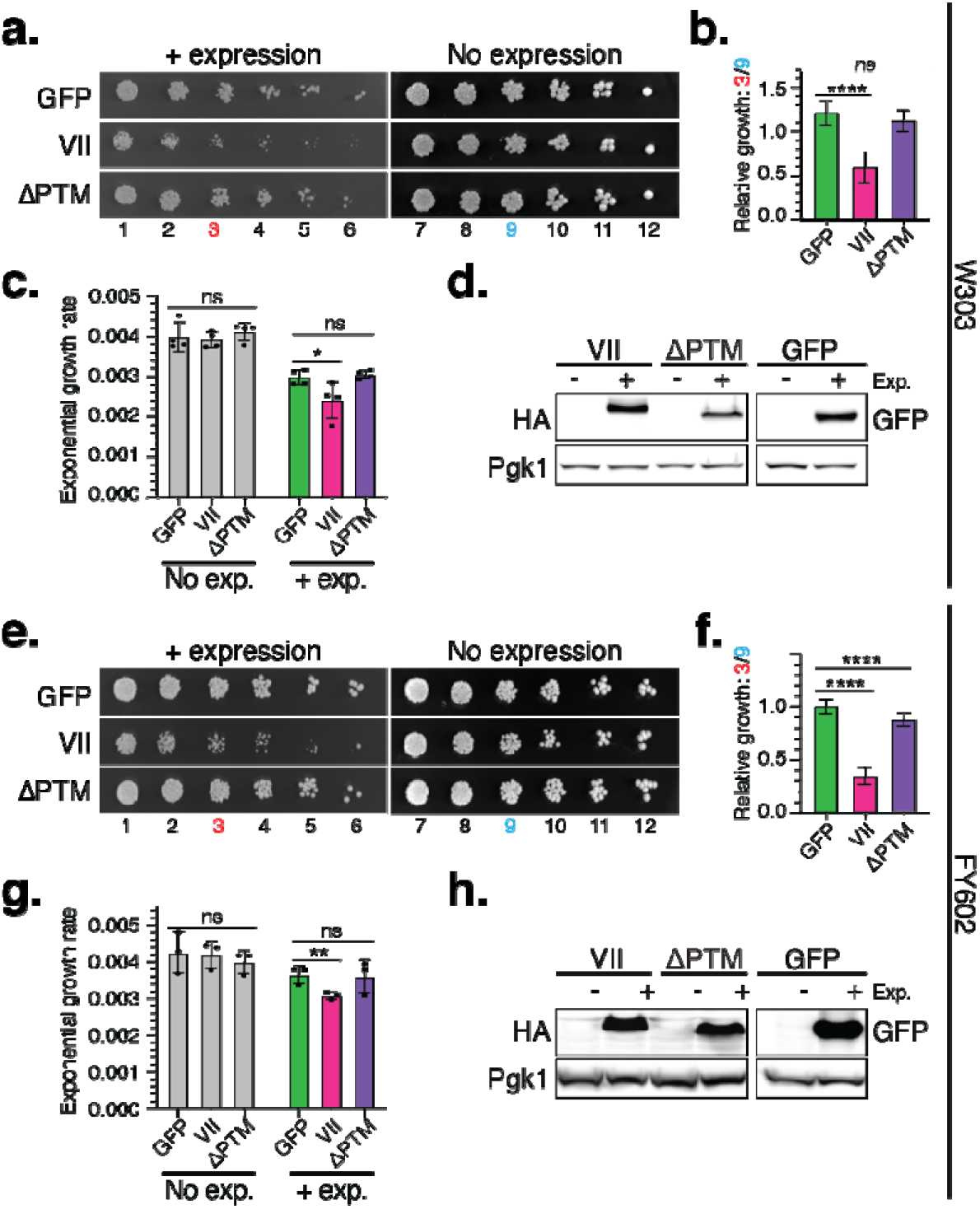
Adenovirus protein VII depletes H1 from chromatin and inhibits growth in budding yeast. **a.** Representative serial dilution assay for growth defects of galactose-induced (+ expression) and dextrose-repressed (No expression) WT haploid W303 cells bearing the inducible protein expression plasmids illustrated in (c). The cells were diluted 1:3 at each step then grown for 3 days at 30°C. **b**. Quantification of WT W303 serial dilution spot assays after 3 days growth at 30°C. The growth intensity for cells with protein expression was measured at dilution spot #3 (red). Growth intensity without expression was measured at spot #9 (blue). The relative growth was calculated as growth intensity at position #3 divided by growth intensity at position #9. Cells expressing VII grew significantly less compared to GFP (*p*<0.0001, one-way ANOVA with multiple comparisons) while cells expressing VII-ΔPTM did not (*p*=0.4069). Error bars represent SD. N=7. **c**. Exponential growth rates measured from WT W303 with the indicated expression plasmids. Using 2-way ANOVA for multiple comparisons, no statistical significance was found among the strains without protein expression (GFP vs. VII *p*=0.9420; GFP vs. VII-ΔPTM *p*=0.7830). Cells expressing VII had a significantly reduced growth rate compared to GFP (*p*=0.0157) while cells expressing VII-ΔPTM did not (*p*=0.9616). Error bars show SD. N=4. **d**. Western blot analysis of WT W303 strains with and without protein expression during exponential growth. Protein VII and VII-ΔPTM are identified with an HA tag and GFP indicates the control strain. Phosphoglycerate kinase (Pgk1) is the loading control. **e**. Representative serial dilution growth assay for growth of WT FY602 with and without GFP, protein VII, or VII VII-ΔPTM expression as described in (d). **f**. Quantification of WT FY602 serial dilution growth assays as described in (e). Cells expressing protein VII grew ~65% less than GFP, which was statistically significant (*p*<0.0001, one-way ANOVA with multiple comparisons), while cells expressing VII-ΔPTM mutant had ~10% less growth than GFP (*p*<0.0001, one-way ANOVA with multiple comparisons). Error bars denote SD. N = 18. **g**. Exponential growth rates measured from WT FY602 with or without protein expression. Using 2-way ANOVA for multiple comparisons, no statistical significance was found among the uninduced strains compared to GFP control (GFP vs. VII *p*=0.9651; GFP vs. VII-ΔPTM *p*=0.3129). Cells expressing protein VII had significantly reduced growth rate compared to GFP (*p*=0.0098) while cells expressing VII-ΔPTM did not (*p*=0.9865). Error bars show SD. N=3. **h**. Western blot analysis of WT FY602 with and without protein expression during log phase growth. Protein VII and VII-ΔPTM are identified with an HA tag, GFP indicates the control strain, and Pgk1 is the loading control.

We found by immunofluorescence microscopy that protein VII is localized to the yeast nucleus (Supplemental Figure 2a), likely as a result of the endogenous nuclear localization signal (NLS) at the N-terminus of protein VII^43^. Therefore, we assessed whether protein VII bound the yeast genome directly by chromatin immunoprecipitation followed by sequencing (ChIP-seq) using an endogenous antibody against protein VII^44^ (Supplemental Figure 2b, Supplemental Figure 3). ChIP-seq showed that protein VII binds throughout the yeast genome. We identified local variability in protein VII binding compared to nucleosomes as determined by total histone H3 ChIP. To test for enrichment of protein VII, we used the MACS2 peak calling algorithm to compare protein VII localization both to the H3 control and to the input control^45^. MACS2 identified very few peaks of protein VII enrichment compared to the input. However, only 9 of those peaks were also called using H3 as the control condition which suggests that they are not bona fide sites of enrichment and that protein VII coats yeast chromatin uniformly just as histone H3 does. Thus, we conclude that protein VII directly binds chromatin throughout the genome. Together these data demonstrate that the effects of protein VII on chromatin in budding yeast are direct and robust, indicating that eukaryotic chromatin is widely vulnerable to protein VII.

### The genetic interactions of protein VII with the yeast homologs of H1, HMGB1, and SET support a linker histone displacement model

Unlike mammals, which have multiple linker H1 and HMGB proteins^10,14^, yeast have primary homologs of linker histone H1, HMGB1 and SET, which are Hho1, Hmo1, and Nap1 respectively^46–51^. To test whether the budding yeast proteins Hho1, Hmo1, and Nap1 interact genetically with protein VII, we introduced the protein VII expression plasmid into strains deleted for each of these genes as well as a double mutant lacking both *HMO1* and *NAP1.* We then assessed whether the deletions affected the growth defect caused by protein VII. We hypothesized that Hmo1 and Nap1, which correspond to the known binding partners HMGB1 and SET respectively, facilitate protein VII dysregulation of chromatin (Figure 2a). Based on this model, we expected that deletion of either the *HMO1* or *NAP1* gene would prevent protein VII from disrupting growth and therefore rescue the growth defects caused by protein VII. Indeed, we found that loss of *HMO1* or *NAP1* during protein VII expression rescued growth compared to WT (Figure 2b). Relative growth with protein VII was significantly increased in *hmo1Δ* compared to WT in the W303 strain (Figure 2c). Likewise, deletion of *NAP1* led to slightly better relative growth compared to WT, though quantification did not reach significance. We also assayed growth with protein VII expression in the same set of gene deletions in the alternate FY602 strain and found that deletion of either *HMO1* or *NAP1* significantly rescued growth (Figure 2e-f). These minor variations in the degree of rescue between the two different strains is likely due to differences in genotype. Taken together, we conclude that both Hmo1 and Nap1 are important for protein VII to cause reduced growth.

**Figure 2.**
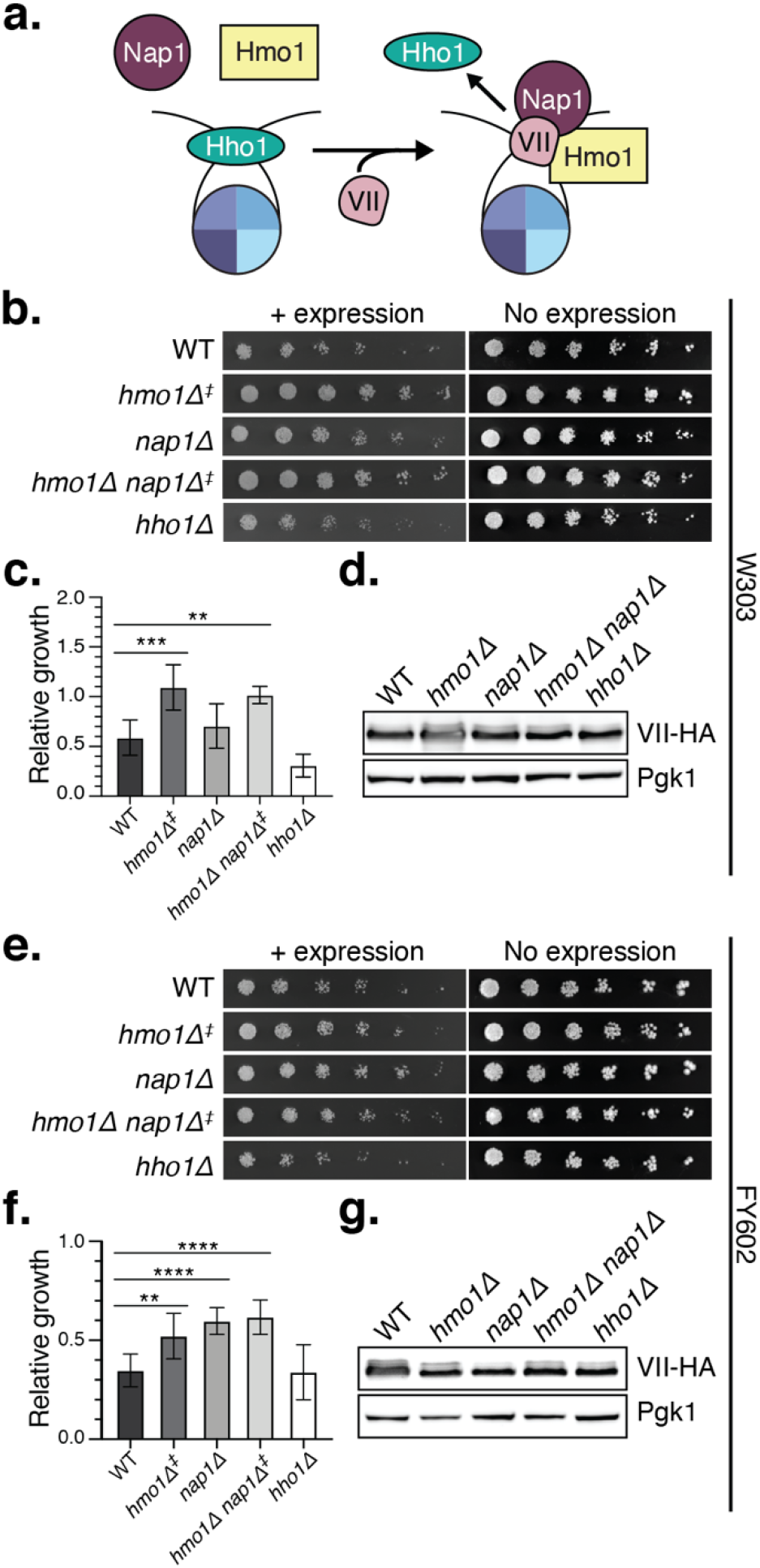
The loss of binding partners Hmo1 or Nap1 rescues protein VII growth defects. **a.** Model for protein VII chromatin disruption in budding yeast. The yeast homologs of HMGB1 and SET, Hmo1 and Nap1 respectively, interact transiently with chromatin, but upon expression of protein VII are recruited to the nucleosome to facilitate protein VII replacement of the linker histone Hho1. **b.** Representative serial dilution assay for growth defects of induced and uninduced W303 cells with protein VII expression plasmid in the indicated gene deletion strains. Due to inherent slowed growth caused by deletion of *HMO1,* spots are shown after 4 days of growth (‡) for *hmo1Δ* and *hmo1Δnap1Δ* strains. All other strains are shown after 3 days of growth. **c.** Quantification of serial dilution growth assays shown in (b). Relative growth was measured as described in Figure 1. Growth density was quantified after 3 days growth for all strains except the *hmo1Δ* and *hmo1Δnap1Δ* strains which were imaged and quantified after 4 days (‡) to compensate for slowed growth caused by the deletion of *HMO1.* Using one-way ANOVA with multiple comparisons against WT, we found that deletion of *HMO1* and deletion of both *HMO1* and *NAP1* in the same strain led to significantly greater relative growth intensity (*p*=0.0009 and *p*=0.0090, respectively). The average growth was slightly higher for the *nap1Δ* strain (*p*=0.7085) and lower for the *hho1Δ* strain (*p*=0.0687) but not significantly. Errors bars show SD. At least three biological replicates were quantified for each strain. **d.** Western blot analysis of induced W303 gene deletion strains shown in (c) with protein VII expression plasmid. Protein VII-HA is shown with Pgk1 loading control. **e.** Representative serial dilution spot assay for growth of FY602 cells with and without induction of protein VII in the indicated deletion strains. Growth after 4 days is shown for the *hmo1Δ* and *hmo1Δnap1Δ* strains (‡) while the other images were taken after 3 days growth. **f.** Quantification of serial dilution growth assay in FY602 deletion strains when protein VII is expressed. Growth density was quantified after 3 days growth for all strains except the *hmo1Δ* and *hmo1Δnap1Δ* strains which imaged and quantified after 4 days (‡). Using one-way ANOVA with multiple comparisons compared to WT, we found that deletion of *HMO1, NAP1,* or both genes together lead to a significant increase in growth intensity (*p*=.0035, *p*<0.0001, *p*<0.0001, respectively). The change in growth intensity when *HHO1* was deleted was not statistically significant (*p*=0.9993). Error bars show SD. At least four biological replicates were quantified for each strain. **g.** Western blot analysis of induced FY602-based strains with protein VII expression plasmid. Protein VII-HA is shown along with the Pgk1 loading control.

We hypothesized that Hmo1 and Nap1 function cooperatively to facilitate protein VII’s disruption of chromatin. As such, removing both factors should have the same effect on the protein VII growth defect as deleting either gene independently. To test this, we assayed growth with protein VII expression in a *hmo1Δnap1Δ* double deletion strain. Consistent with our hypothesis, we found that deletion of both genes rescued the growth defects caused by protein VII to the same degree as a single gene deletion (Figure 2b and e). For both W303 and FY602 strains, the *hmo1Δnap1Δ* deletion strain grew better than WT, but the relative growth was no greater than that of individual deletions of *HMO1* or *NAP1* in the presence of protein VII (Figure 2c and f). Furthermore, we found that protein VII expression levels were largely consistent across strains (Figure 2d and g). The observation that simultaneous deletion of *HMO1* and *NAP1* fails to rescue growth additively supports our hypothesis that these proteins function cooperatively and together give rise to protein VII’s impact on growth.

Based on the finding that protein VII expression reduces H1 chromatin occupancy in human cells (Supplemental Figure 1a) and that protein VII binds linker DNA *in vitro*^32^, we predicted that protein VII and H1 compete for the same binding sites on linker DNA. Thus, we expected that deletion of the yeast H1 homolog, *HHO1,* would allow for more efficient chromatin disruption by protein VII and a corresponding exacerbation of the protein VII growth defect. In both W303 and FY602, we found that without Hho1 growth was reduced compared to WT (Figure 2b and e). While the average relative growth was indeed lower, the reduction was not statistically significant compared to WT (Figure 2c and f). Notably, there were no significant changes in growth upon GFP expression among the strains tested (Supplemental Figure 4). Therefore, we conclude that the changes in growth caused by gene deletion are specific to protein VII expression. In summary, protein VII growth defects are rescued by the deletion of *HMO1, NAP1,* or both factors together while the reduction in growth is exacerbated by *HHO1* deletion. Taken together, these results support our model in which protein VII interacts with Hmo1 and Nap1 to displace Hho1 and disrupt chromatin resulting in growth defects (Figure 2a).

### The human proteins HMGB1 and SET can replace their yeast homologs to facilitate protein VII’s disruption of chromatin in yeast

To determine whether the human factors known to bind protein VII could replace the yeast homologs in our system, we introduced the homologous human proteins into corresponding yeast mutant strains and examined their growth. We assayed the outcome of adding human HMGB1 or SET in the corresponding deletion strains by creating plasmids to co-express protein VII and the target human protein from a bidirectional *GAL1-10* promoter in both W303 and FY602 strains (Figure 3). As observed above, deletion of *HMO1* rescued the protein VII growth defect (Figure 3a-c and g-i). Interestingly, co-expression of HMGB1 with protein VII in the *hmo1\* strain led to reduced growth, which suggests that HMGB1 can replace Hmo1 in facilitating protein VII’s impact on growth. To confirm that these effects on growth were specific to protein VII expression, we tested growth when GFP and HMGB1 were co-expressed in the same set of strains (Supplemental Figure 5a-c and g-i). There was no significant difference in growth when GFP, instead of protein VII, was co-expressed with HMGB1. Thus, we conclude that human HMGB1 can replace yeast Hmo1 to promote the deficient growth caused by protein VII.

**Figure 3.**
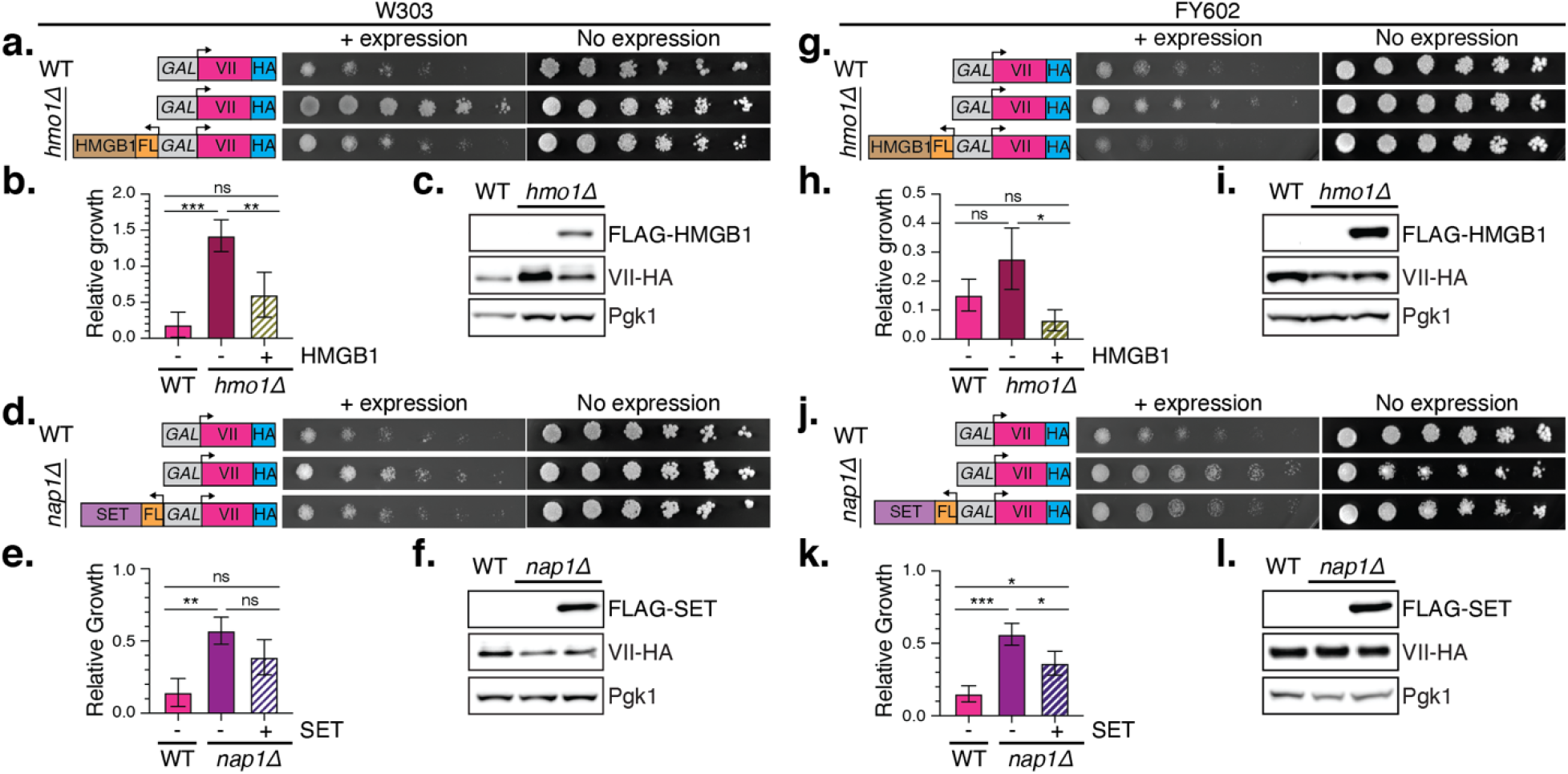
Replacing Hmo1 or Nap1 with homologous human factors exacerbates the protein VII growth defects in yeast. **a.** Serial dilution assay for protein VII growth defects in WT W303 and W303 *hmo1Δ* strain with and without co-expression of human HMGB1. Expression constructs are illustrated to the right. The bidirectional galactose-inducible promoter drives both protein VII and human HMGB1 transcription. Protein VII is HA-tagged and HMGB1 is FLAG-tagged. Cells without protein expression are shown after 3 days growth at 30°C while cells with protein expression are shown after 6 days growth at 30°C to compensate for slowed growth caused by protein VII and the non-preferred energy source, galactose. **b.** Quantification of serial dilution growth assay for strains shown in (a). As in (a), the cells with induced protein expression were quantified after 6 days and cell growth without protein expression was quantified after 3 days growth. A one-way ANOVA with multiple comparisons identified statistically significant differences in growth between WT and *hmo1Δ* strains when VII is expressed (*p*=0.0001). Likewise, growth with protein VII expression in *hmo1Δ* vs. VII coexpression with HMGB1 in the same strain was found to be significantly different (*p*=0.0026). The comparison of growth between WT + VII and *hmo1Δ* + VII + HMGB1 failed to pass the p-value cut-off for significance (*p*=0.068). Error bars represent SD for all growth quantification in this figure. N= 4. **c.** Western blot analysis of WT W303 with protein VII, *hmo1Δ* with VII, and *hmo1Δ* with protein VII and HMGB1 co-expression. HMGB1 is FLAG-tagged, protein VII is HA-tagged, and Pgk1 is the loading control. **d.** Serial dilution assay for protein VII growth defects in WT W303 and W303 *nap1Δ* strain with and without co-expression of human SET as described in (a). The co-expression construct that drives VII and SET from the same bidirectional galactose promoter is illustrated at right. SET is FLAG-tagged, and protein VII is HA-tagged. **e**. Quantification of serial dilution growth assay in WT and *nap1Δ* when protein VII is expressed alone or co-expressed with human SET as performed in (b). A one-way ANOVA with multiple comparisons identified statistically significant differences in growth between WT and *nap1Δ* strains when VII is expressed (*p*=0.0058). When SET was co-expressed with VII in *nap1Δ*, the average growth intensity was reduced compared to VII expression alone in the same strain but not significantly so (*p*=0.0641). No significant difference was found between WT + VII and *nap1Δ* + VII + SET growth (*p*=0.1604). N = 3. **f.** Western blot analysis of WT W303 with protein VII, *nap1Δ* with VII, and *nap1Δ* with protein VII and SET co-expression. SET is FLAG-tagged, protein VII is HA-tagged, and Pgk1 is the loading control. **g.** Serial dilution assay for growth with protein VII expression in WT FY602 and FY602 *hmo1Δ* strain with and without co-expression of human HMGB1 as in (a). **h.** Quantification of serial dilution growth assay in WT FY602 and FY602 *hmo1Δ* when protein VII is expressed alone or co-expressed with human HMGB1 as in (b). While the average growth intensity increased when *HMO1* was deleted compared to WT, the difference was not statistically significant (one-way ANOVA with multiple comparisons, *p*=0.1132). However, replacing Hmo1 with human HMGB1 resulted in a statistically significant reduction in growth (*p*=0.0180). No significant difference in growth was detected when comparing WT and the HMGB1 replacement condition (*p*=0.2982). N = 3. **i.** Western blot analysis of WT FY602 with VII, *hmo1Δ* with VII, and *hmo1Δ* with VII and HMGB1 co-expression. Protein VII is HA-tagged, HMGB1 is FLAG-tagged, and Pgk1 is the loading control. **j.** Serial dilution growth assay for VII expression in WT FY602 and FY602 *nap1Δ* strain with and without co-expression of human SET as described in (d). **k.** Quantification of serial dilution growth assay in WT FY602 and FY602 *nap1Δ* when protein VII is expressed alone or co-expressed with human SET as described in (b). Deletion of *NAP1* rescued the protein VII growth phenotype compared to WT (one-way ANOVA with multiple comparisons, *p*=0.0003). Adding human SET to the *NAP1* deletion strain reduced growth *f*p=0.0240). Replacing Nap1 with SET did not cause the same severity of growth defect as VII expression in WT (*p*=0.0133). **l.** Western blot analysis of WT FY602 with VII, *nap1Δ* with VII, and *nap1Δ* with VII and SET. VII is HA-tagged, SET is FLAG-tagged, and Pgk1 is the loading control.

We performed the same set of assays using human SET to replace yeast Nap1 and obtained similar results (Figure 3d-f and j-l). *NAP1* deletion rescued the protein VII growth defect while expression of human SET in the *nap1Δ* strain restored the protein VII growth deficiency. These observations are consistent with human SET functioning to replace Nap1 in complex with protein VII. When we expressed GFP and SET simultaneously in the *nap1Δ* strain, we found that co-expression did not cause significant changes in growth (Supplemental Figure 5d-f and 5j-l). As was the case with HMGB1, replacing Nap1 with SET altered growth only in the presence of protein VII. Consequently, we conclude that human SET can replace yeast Nap1 to promote protein VII growth defects. Together these data produced by expressing protein VII with human HMGB1 or human SET support our model in which protein VII interacts with these factors to disrupt chromatin.

### Protein VII disrupts the cell cycle in yeast and human cells

To define the cause of slowed growth in the presence of protein VII, we analyzed cell cycle progression by budding analysis in yeast expressing protein VII. Since the yeast budding cycle is correlated to cell cycle progression and can be halted by cell cycle checkpoint activation, a change in the distribution of budding morphologies during log phase growth can indicate that the cells are unable to progress from one phase of the cell cycle to the next^52,53^. We examined the distribution of bud sizes during mid-log phase growth in liquid culture and found significant over-representation of cells with large buds when protein VII was expressed compared to the GFP control (Figure 4a). We next analyzed the budding cycle as the cells transitioned from stationary phase to log phase growth during induction of protein VII expression (Figure 4b, Supplemental Figure 6a). As the cells entered log phase growth, the proportion of large-budded cells increased for both protein VII and GFP expressing cells. However, the cells with protein VII accumulated a significantly greater proportion of large-budded cells which persisted over time. Cells with large buds are likely unable to complete the transition from S phase to G2 or from G2 to M at the same rate as normal cells. Thus, we conclude that protein VII expression disrupts the budding cycle which is consistent with cell cycle progression defects. We also performed budding analysis on the same strains without induction of protein expression and found no significant differences (Supplemental Figure 6b and c), indicating that protein VII specifically leads to the accumulation of large-budded cells. Together these results suggest that the chromatin disruption caused by protein VII in yeast leads to cell cycle dysregulation.

**Figure 4.**
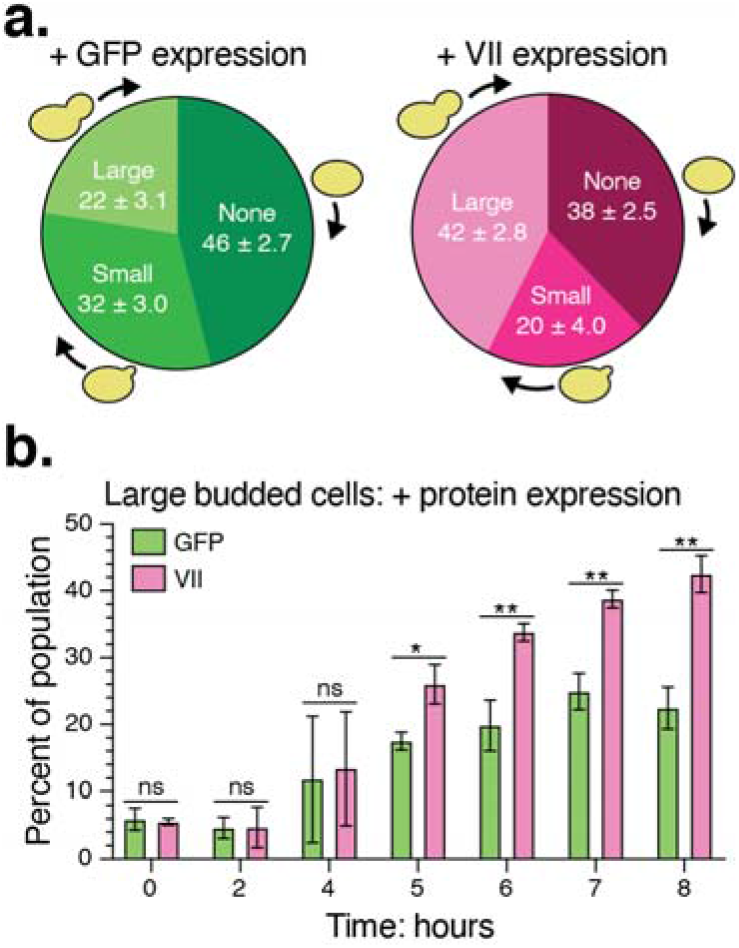
Protein VII dysregulates the yeast budding cycle. **a.** Budding analysis of WT W303 with GFP or protein VII expression mid-log phase growth after 8 hours growth in galactose media. Cells without a bud are in G1 while cells with a small bud are typically in S phase. A large bud roughly indicates G2/M before the cells completes mitosis and cytokinesis to produce a new cell. The percentage of the population with each bud type is shown along with SD. Using multiple unpaired t-tests (two-stage step-up method of Benjamini, Krieger and Yekutieli to control FDR), there was no significant difference in the proportion of unbudded (*p*=0.0172) or small-budded cells (*p*=0.0143). There was significant difference in the proportion of large-budded cells (*p*=0.0012) upon expression of protein VII compared to GFP expressing cells. N=3. **b.** The percentage of large budded cells in the population during protein VII expression (pink) or GFP expression (green) in WT W303. At the beginning of the experiment, the cells in stationary phase (T0) did not express protein VII but were moved to galactose media to initiate protein expression and were grown to mid-log phase (T8). The x-axis indicates time since induction of protein expression and the start of the growth experiment. Unpaired *t*-test *P*-values for each timed sample (T0-T8) from left to right on the x-axis are: 0.727079, 0.962070, 0.837029, 0.010250, 0.003903, 0.001382, and 0.001167. N=3.

To assess the impact of protein VII on human cells, we assayed proliferation and cell cycle progression in human diploid retinal epithelial (RPE-1) cells in the presence of protein VII. Because many adenovirus serotypes can cause conjunctivitis^54^, we chose these cells as biologically relevant for adenovirus infection. We expressed GFP-tagged protein VII or a GFP control by a recombinant viral vector then measured cell proliferation and cell cycle progression by flow cytometry to detect DNA content^55^. We found that cells expressing protein VII stopped proliferating immediately, similarly to control cells arrested by treatment with a CDK-1 inhibitor, RO-3306^56^ (Figure 5a; Supplemental Figure 7a and b). Given that more than 85% of the protein VII-GFP cells excluded trypan blue stain, we concluded that this loss of proliferation was not caused by cell death (Supplemental Figure 7c). Rather, we found that protein VII-GFP expression led to an increase in G2/M DNA-content cells compared to the control (Figure 5b). Since flow cytometry analysis for DNA content does not distinguish between G2 and mitotic cells, we examined phosphorylation of histone H3 at serine 10 (H3S10ph) as a marker of mitotic chromosomes^57–59^. Protein VII-GFP expressing cells had dramatically higher H3S10ph levels compared to the other conditions (Figure 5c). The amount of H3S10ph increased relative to the pre-treatment sample and remained high over time suggesting that the protein VII-GFP cells enter mitosis but are unable to complete it. In agreement with this conclusion, we observed more binucleate cells in the population of protein VII-GFP cells (data not shown). In contrast, the GFP and no treatment samples had a steady decline in H3S10ph which likely reflected the reduction in G2/M cells detected by flow cytometry. The RO-3306-treated cells had nearly undetectable levels of H3S10ph which is consistent with Cdk1-dependent activation of aurora A/B kinases during the G2/M transition^60^. Together, these results indicate that protein VII slows the cell cycle in diploid human cells leading to an enrichment of mitotic cells, which echoes our observations in yeast. Hence, we conclude that expression of protein VII impairs progression of the cell cycle through mitosis.

**Figure 5.**
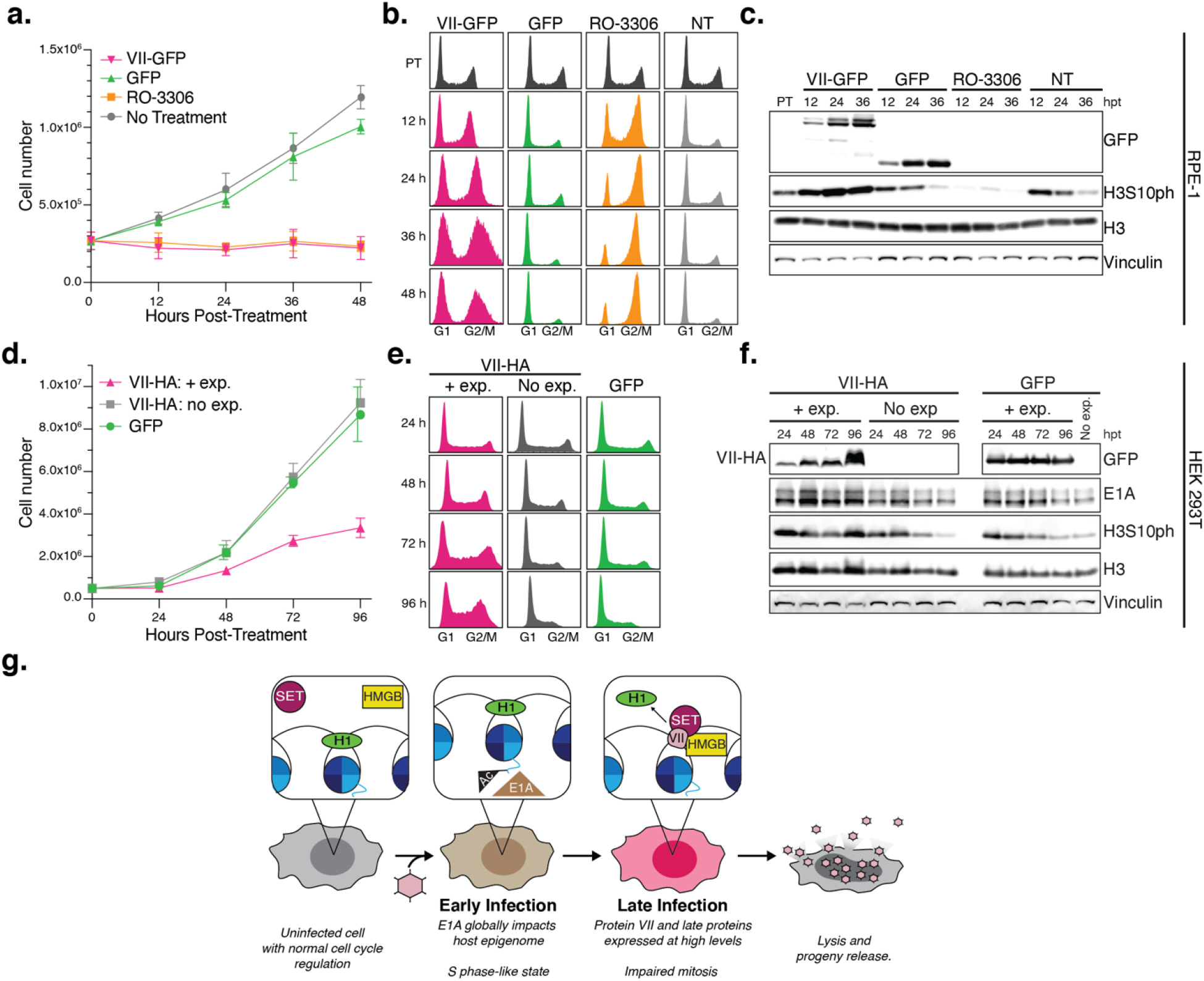
Protein VII blocks proliferation and impedes cell cycle progression in human cells. **a.** Diploid RPE-1 cell proliferation for 48 hours after treatment. The cells were transduced with recombinant adenovirus to express GFP-tagged protein VII or a GFP control. Protein VII-GFP recombinant adenovirus MOI = 50. GFP control recombinant adenovirus MOI = 50. 10 μM RO-3306 CDK-1 inhibitor treatment is a control for inhibition of proliferation and G2 arrest. At 48 hours, there was a significant difference in the total number of protein VII-GFP expressing cells compared to the no treatment control (*p*<0.0001, one-way ANOVA with Tukey’s multiple comparisons test) and to the GFP control (*p*<0.0001). There was no difference between the cells expressing protein VII-GFP and RO-3306 treated cells (*p*=0.9933). Error bars show standard deviation. N = 4. **b.** Cell cycle analysis by flow cytometry of transduced RPE-1 cells shown in (a). Cells were fixed and stained with propidium iodide to assess DNA content. DNA content is plotted on the x-axis, with G1 and G2/M populations labelled, and the y-axis represents a histogram of the cell population. The y-axis is normalized to fill the entire axis independently for each sample. Since nearly 100% of transduced cells were positive for GFP expression, cells were not gated on GFP positive cells. The G1 and G2/M populations correspond to cells with 2N and 4N DNA content respectively. **c.** Western blot analysis of transduced RPE-1 cells shown in (a). Samples labeled “NT” are the no treatment control. “PT” indicates the pre-treatment sample and “hpt” shows the time post-treatment in hours. The GFP blot shows the relative protein VII-GFP and GFP expression. H3S10ph indicates mitotic chromosomes. Total histone H3 and vinculin loading controls are also shown. **d.** HEK 293T proliferation with and without protein VII-HA expression. The protein VII-HA + expression and GFP cells were induced to express the target proteins under a doxycycline inducible promoter for 96 hours. The protein VII-HA no expression cells were not treated with doxycycline. At 96 hours post-treatment, there were significantly fewer cells expressing protein VII-HA compared to the GFP control and the untreated protein VII-HA cells (one-way ANOVA with multiple comparisons; *p*=0.0016 and *p*=0.0009 respectively). There was no significant difference in cell number between the GFP control and untreated protein VII-HA cells (*p*=0.7841). Error bars represent SD. N = 3. **e.** Flow cytometry cell cycle analysis of HEK 293T cells shown in (d) using DNA content to assess cell cycle progression as described (b). Since more than 90% of the doxycycline treated GFP cells were positive for GFP expression, the cells were not gated on positive expression. DNA content is shown on the x-axis and G1 and G2/M peaks are indicated although the ploidy of HEK 293T cells varies within the population of cells. The y-axis histogram is normalized to fit the entire G1 peak for each sample. **f.** Western blot analysis of HEK 293T cells shown in (d). HA- and GFP-probed blots indicate expression of target proteins in doxycycline-treated samples. E1A indicates the presence of adenovirus early expressed E1A protein which contributes to the transformation of HEK 293T cells. Multiple E1A isoforms are resolved as multiple bands. H3S10ph indicates mitotic chromatin. Total histone H3 and vinculin are loading controls. **g.** Model for protein VII on host chromatin during infection. An uninfected cell has normal cell cycle regulation. Upon infection, the early-expressed adenovirus genes E1A and E1B bypass cell cycle checkpoints and push the cell into an S phase-like gene expression program. E1A acts primarily through redistributing acetylation of histone H3K18 globally. Late during infection, abundant protein VII recruits HMGB1 and SET to displace the linker histone from host chromatin. Protein VII chromatin disruption impedes mitosis and therefore cytokinesis, culminating in cell lysis and maximizing the release of viral progeny.

### Protein VII blocks proliferation of HEK 293 cells

During adenovirus infection, protein VII is expressed from the L2 late transcriptional unit^61,62^. The finding that ectopically expressed protein VII is a potent inhibitor of proliferation and cell cycle progression in human cells is in stark contrast to the well-studied effects of the early adenovirus genes E1A and E1B, which direct the bypass of cell cycle checkpoints and promote an S phase-like gene expression state^31,63^. This E1A/E1B gene expression program is the primary driver of transformation in the ubiquitous human cell line HEK 29 3^64,65^. Consequently, we asked whether protein VII could effectively override the pro-proliferation effects of E1A.

To test whether protein VII can abrogate the effects of E1A on proliferation and cell cycle regulation, we generated an E1A/E1B-transformed cell line to express HA-tagged protein VII under a doxycycline-inducible promoter from a single locus using the HILO-RMCE system^66^. After induction of protein VII expression, growth slowed significantly over time compared to the GFP control cells (Figure 5d, Supplemental Figure 7d and e). The loss of proliferation in the inducible protein VII 293 cells occurred more gradually than in the RPE-1 cells likely due to the lower levels of protein VII expressed upon initial induction compared to transduction (Figure 5f).

Like the RPE-1 cells, protein VII expression decreased the number of G1 cells and increased the proportion of G2/M cells (Figure 5e) resulting in an accumulation of H3S10ph (Figure 5f). The growth impairment was not caused by widespread cell death (Supplemental Figure 7f) nor by loss of E1A expression (Figure 5f). Thus, we conclude that protein VII is able to counteract the effects of E1A and halt cell proliferation. Together these data suggest that during adenovirus infection, the late-expressed protein VII can counteract the effects of E1A on cell cycle regulation and prevent cell division. This newly uncovered role for protein VII suggests that disruption of the cell cycle late in infection may be as critical to viral replication as cell cycle disruption during the earliest phases of infection.

## Discussion

For nuclear-replicating viruses such as adenovirus, host chromatin is both an impediment to and a bountiful resource for ensuring successful replication. The histone-like protein VII is an important, yet largely mysterious, component of adenovirus’ armament for re-purposing host chromatin resources. Previous work showed that protein VII directly interacts with protein SET^67,68^ and HMGB1 and recruits them to chromatin during infection^32^. Proteomic analysis of chromatin in the presence of protein VII revealed that linker histones are depleted while SET and HMGBs are enriched, suggesting that protein VII may co-opt these host factors to disrupt linker histone occupancy^37,69,70^. As such, we hypothesized that protein VII exploits the antagonism between H1 and HMGB1 to undermine chromatin structure and maximize viral success. Here, we coupled budding yeast genetics and human cell experiments to test our model of linker histone displacement and chromatin disruption via SET and HMGB proteins.

The expression of protein VII elicited a significant growth defect in two separate lab yeast strains, demonstrating the robustness of protein VII’s chromatin perturbation. Protein VII expression led to smaller colony size, indicating slowed growth, while fewer colonies grew on the solid media suggesting a loss of viability or proliferation (Figure 1). This is consistent with the large-budded cell phenotype observed during exponential growth (Figure 4) and suggests that cell cycle checkpoint activation leads to cell death or reduced growth rate. As a control, we expressed the VII-ΔPTM protein which, in human cells, cannot be post-translationally modified and therefore no longer localizes to chromatin. In yeast, expressing VII-ΔPTM did not cause growth defects suggesting that these sites are also critical for protein VII’s disruption of chromatin in yeast. It is possible that protein VII is modified in yeast and that these modifications regulate protein VII’s interaction with and impact on chromatin. It may also be the case that structural changes caused by the mutations render protein VII unable to impact chromatin despite stable expression. While protein VII bound chromatin across the yeast genome and thus was not significantly enriched at specific loci, the genomic profile is distinct from that of total H3 (Supplemental Figures 2 and 3). Further work focused on protein VII occupancy will define the specificity of protein VII binding genome wide.

Deleting *HMO1* or *NAP1,* the yeast homologs for HMGB1 and the histone chaperone protein SET, rescued the growth defects caused by protein VII expression while deletion of the linker histone, *HHO1,* exacerbated this phenotype. Interestingly, *HMO1* deletion had a more profound effect in W303 while *NAP1* deletion led to greater rescue in FY602 (Figure 2). Furthermore, when both factors were deleted, the magnitude of rescued growth matched that of a single gene deletion. For example, in W303, the relative growth of the *hmo1Δnap1Δ* strain was roughly equal to that of the *hmo1Δ* strain. The lack of an additive rescue indicates that these factors function in the same genetic pathway with respect to protein VII. However, the strain-specific results suggest that the relationship between Hmo1 and protein VII has a greater impact in W303, while protein VII’s effect is mediated more by Nap1 in FY602. These unique strain relationships are also evident upon re-introduction of the human homologs. For both strains, expressing the homologous human factors when protein VII was present rescued the growth phenotype in the corresponding deletion strains (Figure 3). However, expression of HMGB1 in the W303 *hmo1Δ* strain had a greater impact on growth than the expression of protein SET in the *nap1Δ* strain. We observed the opposite relationship in FY602. Given that W303 has nearly ~9500 single nucleotide variations affecting nearly 12% of all genes when compared to FY602^71^, it is likely that this considerable variation between the two strains gives rise to the different apparent preference for Hmo1 or Nap1. In neither case, however, did expression of the human factor completely rescue the protein VII growth defect suggesting that other factors contribute to protein VII’s disruption of chromatin.

Consistent with our model that protein VII and H1 compete for the same binding sites, deletion of the linker histone exacerbated the effects of protein VII on yeast growth (Figure 2). As such, H1 may protect chromatin from protein VII invasion. Based on the protein VII ChIP-seq from yeast chromatin (Supplemental Figures 2 and 3), we propose that protein VII binds human chromatin ubiquitously, without preference for DNA sequence or chromatin state. However, locations that have higher H1 occupancy may be resistant to protein VII binding, and thus protein VII requires the action of protein SET and HMGB proteins to facilitate its chromatin invasion. This would explain why the effects *of HHO1* deletion in yeast are relatively slight with respect to protein VII growth impairment. Compared to human cells, yeast have very little heterochromatin which is silenced by means other than linker H1 binding^72^. Therefore, a large portion of the yeast genome is vulnerable to protein VII invasion, even without *HHO1* deletion, compared to the human genome which is thought to be comprised of ~25% heterochromatin^73^. Furthermore, the post-translational modifications of H1 proteins, which have not been well characterized, may regulate H1 localization and antagonism of HMG proteins. Given that post-translational modification of protein VII impacts its relationship with chromatin, it is possible that protein VII competes for these H1-modifying enzymes to cause chromatin dysregulation in yet another dimension.

Similar to our observations in yeast, protein VII resulted in a dramatic loss of proliferation and cell cycle arrest in diploid RPE-1 cells (Figure 5). The block of cell growth and shift in cell cycle profiles occurred rapidly, within 12 hours of protein VII expression. However, the block of growth took longer in the HEK 293 cells likely due to lower levels of protein VII expression driven from a single copy of the gene as opposed to the multiple copies introduced to each RPE-1 cell by transduction. While the most profound effects on cell cycle progression seemed to occur during mitosis, we also observed an increased S phase population in both human cell types. This build-up of S phase cells suggests that protein VII’s interruption of cell cycle progression can occur at multiple points.

In HEK 293 cells, E1A promotes proliferation and transforms the cells by re-wiring their basal epigenetic state. This occurs through redistributing H3 acetylation which then alters global gene expression^26,74^. E1A most significantly upregulates acetylation of H3K18 to reprogram transcription^1^. In contrast, protein VII expression does not cause significant changes to H3K18ac or most other histone marks^32^. This suggests that the mechanism of protein VII’s block of proliferation in HEK 293 cells is not through changes to histone modifications. Likewise, although protein VII and E1A have been shown to directly interact^75^, the effects of protein VII on proliferation and cell cycle regulation occur through means independent of this interaction given that these effects are observed in cells without E1A expression (Figure 5) and in budding yeast (Figure 1).

It is remarkable that, although budding yeast and humans are separated by one billion years of divergent evolution and yeast have not encountered adenovirus infection during that evolutionary history, adenovirus protein VII disrupts yeast chromatin just as it does human chromatin. This finding underscores both the conservation of chromatin biology across eukaryotes as well as the robust and catastrophic effects of protein VII on the most basic functions of chromatin. Our study, as well as others^76^, demonstrates the power of yeast to study the effects of histone-like proteins on chromatin. In light of this new role for protein VII in undermining cell cycle progression, we propose that protein VII expression late in the infection cycle maximizes progeny production by counterbalancing checkpoint bypass and transcriptional reprogramming initiated by the early-expressed factors E1A and E1B (Figure 5g). These earliest expressed adenovirus proteins induce a “viral S phase”^77,78^ while a complement of other early adenovirus proteins inactivate the host DNA damage response^79^. Since both the intra-S phase and G2 host cell cycle checkpoints rely on sensing DNA damage to stall cell cycle progression, it is conceivable that, as a consequence of this un-checked “viral S phase”, an infected cell could initiate mitosis and cytokinesis before viral progeny assembly and release. In this scenario, breakdown of the nucleus would disrupt progeny assembly and cell division would redirect nuclear resources away from viral replication resulting in reduced progeny. Given the effects of protein VII on mitosis, we propose that protein VII prevents an infected cell from initiating cytokinesis, therefore maximizing virion production. Studies of several mutant adenoviruses provide indirect evidence that late viral proteins can prevent the onset of mitosis. For example, cells synchronized in early G1 then infected with mutant adenovirus lacking E1B-55k and E4orf3 are predisposed to arrest in a mitosis-like state late in infection^78^. It’s likely that mutant viruses do not highly express late proteins, including protein VII. Therefore, given our observation that protein VII impedes mitosis without the need for other early adenovirus proteins, it will be informative to test the contribution of protein VII and its binding partners to cell cycle progression during infection.

In sum, we show that adenovirus protein VII disrupts chromatin by exploiting H1-HMGB antagonism and, as a result, impairs cell cycle progression. Our findings in yeast and human cells reveal a highly conserved chromatin vulnerability that adenovirus exploits, underpinning the evolutionary conservation of H1-HMGB antagonism. HMGB1 has long been the focus of studies of host inflammatory responses^80,81^, but its intranuclear role during infection is also intimately linked to viral infection^32^. Intriguingly, HMGB1 was recently found to be required for SARS-CoV-2 infection^82^. Although the mechanism is not known, preliminary evidence suggests that the interaction of HMGB1 with chromatin underlies this dependency, which hints that the novel vulnerability we expose in H1-HMGB1 antagonism may be exploited by a wide range of intracellular pathogens.

## Acknowledgements

We thank members of the Avgousti lab, M. Emerman, M. Weitzman, E. Hatch, S. Parkhurst and T. Tsukiyama for insightful comments on the manuscript. We also thank H. Wodrich for generously providing anti-VII antibodies and D. Curiel for sharing recombinant adenoviruses. We also thank N. Donadio for technical help, the Hatch lab for assistance with cell cycle dynamics, the Brewer/Raghuraman lab for providing yeast strains, L. Howe for providing yeast strains and constructs, and the Tsukiyama lab for invaluable advice, reagents and insight for yeast and chromatin experiments. We also thank the Fred Hutch Cellular Imaging Shared Resource for assistance with microscopy and image analysis. This research was supported by the Cellular Imaging Shared Resource (CISR) and Genomics Core Facility of the Fred Hutch/University of Washington Cancer Consortium (P30 CA015704). We also thank the Bioinformatics and Flow Cytometry Facilities at Fred Hutchinson Cancer Research Center for their assistance. The research presented here was supported in part by start-up funds from Fred Hutchinson Cancer Research Center and NIH funding to DCA (R35-GM133441), KLL (T32-CA009657), and HCL (T32-AI083203).

## Author Contributions

Conceptualization, K.L.L, D.C.A.; Methodology, K.L.L, D.C.A.; Formal Analysis, K.L.L., M.B., M.R.D.; Investigation, K.L.L., M.B., M.R.D., H.C.L., D.C.A.; Resources, K.L.L., M.B., M.R.D., H.C.L., D.C.A.; Data Curation, K.L.L., M.B., M.R.D.; Writing – Original draft, K.L.L and D.C.A.; Writing – Review & Editing, K.L.L., M.B., M.R.D., H.C.L., D.C.A.; Visualization, K.L.L., M.B., M.R.D.; Supervision, K.L.L, D.C.A.; Funding Acquisition, K.L.L, D.C.A.

## Materials and Methods

### Yeast Expression Plasmids

We used a standard *GAL1* promoter plasmid and inserted a protein VII sequence codon optimized using the IDT algorithm for expression in *S. cerevisiae.* To do so, we synthesized gBlock Gene Fragments (Integrated DNA Technologies) corresponding to mature protein VII from human adenovirus type 5 and a version of the same protein coded for alanine substitutions at five residues previously found to be post-translationally modified^32^. Both versions of protein VII were encoded with three consecutive HA epitope tags at the C terminus. These protein VII DNA fragments, as well as yeast codon-optimized GFP sequence from the plasmid PFA6A-GFP-KanMx^83^, were PCR amplified to have 3’ and 5’ SpeI and HindIII restriction sites respectively. The amplified sequences were then cloned and confirmed by Sanger sequencing across both junctions. To generate the co-expression plasmids with either WT protein VII or GFP, the *GAL1* promoter was first replaced with the full-length bidirectional *GAL1-10* that was PCR amplified from the FY602 strain and engineered with XbaI and SacI restriction sites. We confirmed successful replacement with the bidirectional *GAL1-10* promoter by Sanger sequencing. Next, we PCR amplified human protein SET or human HMGB1 DNA fragments (codon optimized for yeast) containing SacI restriction sites at both ends of the fragments from plasmids synthesized by GeneWiz. These fragments were then cloned opposite either GFP or protein VII allowing for co-expression of either gene with human protein SET or HMGB1.

### Yeast Strain Construction and Maintenance

We obtained WT FY602 from from Dr. Leann Howe at the University of British Columbia^76^ and introduced gene deletions for *HMO1, HHO1,* and *NAP1* using one-step homologous recombination gene replacement as in reference ^84^. The *HIS3* gene used for replacement of *HMO1* was amplified from pRS413^85^ and the *KanMx6* gene used for replacement of *HHO1* or *NAP1* was amplified from pUG6^86^. WT W303 was obtained from the Brewer-Raghuraman lab and all W303-derived deletion strains were obtained from the Tsukiyama lab. The linear deletion fragments or plasmids were introduced by standard lithium acetate transformation^87^ and growth on selective media. Transformed colonies were streaked out for single isolates on selective media and candidate gene deletions were confirmed by diagnostic PCR to confirm integration of the selection gene as well as Sanger sequencing of the insertion junctions. For strains obtained from other labs, putative gene deletions were confirmed by diagnostic PCR to confirm integration of the selection gene. Expression plasmids were transformed into relevant strains every four weeks. Colonies were streaked onto selective media, grown at 30°C until single colonies formed, and then stored at 4°C. Strains containing the expression plasmids were maintained in selective media for all pre-growth steps.

### Yeast Media

Yeast media was prepared using ultrapure water (Millipore Sigma Synergy UV Remote Water Purification System, #SYNSVR0WW) and sterilized with 15-minute liquid autoclave cycle. Standard rich media (YEP) was prepared with bactopeptone (Fisher # DF0118-17-0), yeast extract (Fisher #BP1422-500), and 2% dextrose (Fisher Bioreagents #D16-500) with 2.4% agar (Fisher Bioreagents #BP1423-500) for agar plates. Kanamycin was added to 400 mg/mL in YEPD plates for selection. Complete media without uracil (SC-ura) or histidine (SC-his) was purchased from Sunrise Science Products (#1306-030 and #1303-030) and prepared per the manufacturer’s instructions with yeast nitrogen base (#1501-250) and with 2.4% agar for solid media plates. To derepress the galactose-promoter, cells were grown in SC-ura + 2% raffinose media (Fisher #AC195675000). The induce protein expression, cells were grown in SC-ura + 2% galactose media (Fisher #BP656-500).

### Yeast Growth Assays and Quantification

For serial dilution growth assays, plasmid-bearing strains were pre-grown for 24 hours in SC-ura + 2% dextrose liquid media at 30°C with agitation. Cultures were sonicated at 35% amplitude for 7 seconds using a Fisher Scientific FB50 Sonic Dismembrator to disaggregate the cells before dilution. Cells were diluted in ultrapure water and counted by hemocytometer, then six 1:3 serial dilutions were prepared in a 96-well plate such that the most dilute cell mixture contained 3 cells/uL. After vigorous trituration, 5 uL of each solution was placed onto SC-ura + 2% dextrose agar media to repress protein expression or SC-ura + 2% galactose agar media to induce protein expression. The plates were dried at room temperature and then incubated at 30°C for six days. Each day, the plates were imaged on a BioRad ChemiDoc MP Imaging System using the “Ponceau S” setting and 0.5s exposure time. The images were processed using Adobe Creative Cloud Photoshop and Illustrator. The serial dilution assays were quantified using BioRad Image Lab v6.1 software. Using the Volume Tools analysis package, a box with the same pixel dimensions was drawn around the third position in the dilution series (see Figure 1) for both the dextrose- and galactose-grown cells and the signal intensity values were measured for each position independently. The local subtraction method was used to remove background signal. To determine the relative growth value, the signal intensity of grown on galactose media was divided by the signal intensity from dextrose media growth. For WT, *nap1Δ* and *hho1Δ* strains with the single protein expression plasmids, which all had the same growth rate, the relative growth values were calculated from the images taken after three days growth at 30°C.

To compensate for slowed growth caused by Hmo1 deletion, the relative growth for *hmo1Δ* and *hmo1Δnap1Δ* strains was measured after four days growth. For strains with the co-expression plasmids, which had slightly different growth rates, relative growth was calculated after five days growth during GFP expression or six days growth during protein VII expression. Quantification of the serial dilution assays represent at least three independent replicate experiments. Statistical tests were carried out as described in the figure legends using GraphPad Prism v9.0 software.

For the exponential growth analysis, strains were pre-grown for 24 hours in SC-ura + 2% dextrose liquid media at 30°C with agitation, then scaled up to a larger culture volume in SC-ura + 2% raffinose liquid media and grown for another 24 hours. To start the exponential growth experiments, the raffinose-grown cultures were diluted to OD_660_ of 0.15 in SC-ura + 2% dextrose or SC-ura + 2% galactose and incubated at 30°C with agitation. The OD_660_ was measured every hour and graphs were generated in real time. Western blot samples for protein expression confirmation were collected at 4-, 6-, 8- and 10-hours post-induction for the first experimental replicate and at 6 hours post-induction for follow-up experiments. To determine exponential growth rate, the OD_660_ was graphed on a log-10 scale and an exponential growth equation with least squares fit was fit to at least five datapoints using GraphPad Prism v9.0 software. The bar charts represent the average exponential growth value (k) from at least three biological replicate experiments. Statistical tests are described in the figure legends and were performed with GraphPad Prism v9.0 software.

### Yeast Western Blotting

To confirm protein expression, western blot samples were recovered after six hours induction or repression during exponential growth. To equalize the number of cells in each sample, culture volume containing 2 OD_660_ of cells was centrifuged to pellet the cells. The cells were washed once with filtered purified water, pelleted again, and the dry pellets were stored at −80°C until processing. To prepare whole cell lysate, the cell pellet was resuspended in 100 uL water then 100 uL 0.2M NaOH was added and cells were incubated at room temperature for five minutes^88^. The cells were pelleted, then resuspended in SUMEB loading buffer (1% SDS, 8 M Urea, 10 mM MOPS, pH 6.8, 10 mM EDTA, 0.02% bromophenol blue) with protease inhibitor cocktail (MilliPore Sigma #11836170001) and 5% 2-mercaptoethanol followed by heating at 65°C for 10 minutes. Samples were loaded onto 15% SDS-PAGE gels and run via standard methods in NuPAGE MOPS SDS running buffer (Invitrogen #NP0001). Gels were transferred (Transfer Buffer: 25 mM Tris Base, 100 mM glycine, 20% methanol) onto nitrocellulose membrane and Ponceau stained, blocked with 5% milk for 30 minutes, and stained with primary antibody overnight. Primary antibodies and dilutions used were: α-GFP (1:2500; Abcam #290), α-HA (1:500; BioLegend #MMS-101R), α-PGK1 (1:5000; Abcam #113687), and α-H3 (1:2500; Abcam #1791). After standard washing, blots were probed with either α-mouse (1:5000; Jackson ImmunoResearch #115-035-003) or α-rabbit (1:5000; Jackson ImmunoResearch #111-035-045) HRP-conjugated secondary antibody and developed with ECL per the manufacturer’s instructions (BioRad Clarity ECL #1705061). Images were acquired using the BioRad ChemiDoc MP Imaging System.

### Yeast Immunofluorescence

Cultures were pre-grown in liquid SC-ura + 2% raffinose media as previously described, then scaled to 50 mL culture volume at an OD660 of 0.3 in SC-SC-ura + 2% dextrose or SC-ura + 2% galactose liquid media. Cultures were incubated for 4 hours at 30°C with agitation. After induction, culture OD_660_ was measured such that equivalent numbers of cells could be collected by centrifugation. Cells were sonicated for 10 seconds at 50% duty cycle, output 2 using a Branson Sonifier 250 Model SSE-1 probe sonicator (#510-294-301). The cells were pelleted by centrifugation at 3000 rpm for 5 minutes, washed twice in water, resuspended in 1 mL 4% PFA diluted PBS, and incubated for 1 hour at room temperature on a nutating platform mixer to fix the cells. Cells were pelleted by centrifugation as before, washed and then resuspended in 1mL Wash Buffer (0.1 M KH_2_PO_4_ in 1.2M sorbitol solution). Cells were counted by hemocytometer and 40 million cells were moved to a fresh microtube and brought up to 200 uL total volume in Wash Buffer. Zymolyase (10 mg/mL Zymo Research #ZE1005) and 2-mercaptoethanol was added to the cells which were incubated for 50 minutes at 30°C to spheroplast the cells. Cells were washed twice by pelleting then resuspension in 1 mL Wash Buffer. Cells were then resuspended in 200 uL 3% BSA solution in PBS and incubated at room temperature with agitation for 30 min. Cells then incubated in primary antibody solution containing α-HA (1:400; Abcam #9110) and α-PGK1 (1:400; Abcam #113687) antibodies in the 3% BSA solution, then incubated for 1 hour with rocking at room temperature out of direct light. Cells were washed thrice in 3% BSA solution, then suspended in 200 uL secondary antibody solution containing α-rabbit Alexa Fluor 488 (1:300; Invitrogen #A11008), α-mouse Alexa Fluor 555 (1:300, Invitrogen #A28180), and 2 ug/mL DAPI (Fisher #50-874-10001). Cells were washed once in 3% BSA solution, three times in PBS, and resuspended in 50 uL PBS. Cells were counted by hemocytometer and diluted to a final concentration of 500,000 cells per uL in PBS. 2 uL of cell solution was mixed with Invitrogen ProLong Gold Antifade (#P36934) and mounted on poly-lysine slides. Slides were visualized with a Zeiss 780 LSM confocal microscope, maximum intensity projection images were processed in FIJI and assembled using Adobe Photoshop and Illustrator.

### Yeast Chromatin Purification and Native Chromatin Immunoprecipitation (ChIP-seq)

After 6 hours galactose induction during log-phase growth in liquid culture, 500 mL culture volume of cells was collected, nuclei purified, chromatin isolated, and native ChIP performed as previously described^89^. The ChIP protocol was performed using α-H3 (Abcam #1791) and α-protein VII^44^ antibodies and Invitrogen Protein G Dynabeads for Immunoprecipitation (ThermoFisher catalog #10004D). Libraries were prepared from the ChIP samples as well as a 10% input, following manufacturer recommendations, with KAPA Biosystems Unique Dual-Indexed Adapter Kit (#KK8727), Hyper Prep Kit (#KK8504), and Beckman Coulter AMPure XP Beads (#A63880) for library cleanup and size selection. Library concentrations were measured with Invitrogen Qubit dsDNA HS Assay Kit (#Q32851), then analyzed with Agilent High Sensitivity D5000 ScreenTape System, and pooled. Libraries were sequenced with 50-bp paired-end reads on an Illumina NovaSeq 6000 SP sequencer at the Fred Hutch Genomics Core Facility. Reads were aligned to the Saccer3 yeast genome assembly (GenBank accession GCA_000146045.2) using Bowtie2. Read coverage was assessed using bamCoverage package from DeepTools2.0. Reads aligning uniquely to the yeast genome were used to identify protein VII peaks using the MACS2 peak caller. Peaks designated as “narrow” and bedGraph tracks showing read coverage for the input, H3, and protein VII ChIP samples for one replicate are shown in Supplemental Figures 2 and 3. ChIP-seq was performed in duplicate. All ChIP-seq sequencing data are available at GEO accession #GSE164684.

### Yeast Budding Index

FY602 strains were pre-grown in raffinose and transferred to SC-ura + 2% dextrose or SC-ura + 2% galactose liquid media as during exponential growth analysis described above. Samples were collected every 1-2 hours during growth/induction. 500 uL of each culture was pelleted, washed once in MilliQ water, resuspended in 70% ethanol, and fixed overnight at 4°C. Samples were pelleted, washed once with 50 mM sodium citrate, resuspended and diluted in the same solution. Cells were sonicated as described above and then loaded onto a hemocytometer counting chamber. 120-170 cells per sample were classified as “unbudded”, “small-budded” (bud < 50% the size of the mother cell), “large-budded” (bud > 50% the size of the mother cell), or “irregular” (multiple buds or other irregularities). Data were graphed and statistical analyses performed as described in the figure using GraphPad Prism v9.0 software.

### Human Cell Line Generation and Maintenance

hTERT-immortalized RPE-1 cells were obtained from Dr. Emily Hatch at Fred Hutchinson Cancer Research Center and were grown in 1:1 DMEM/F12 media (Thermo Fisher #11330032) with 10% FBS (Sigma #FO926-500ML; Lot #18C539), 1% penicillin-streptomycin (Fisher #15-140-122), and 0.01% hygromycin (Sigma #H3274-50MG). HEK293T HILO acceptor cells were obtained from Dr. Eugene Makeyev. Cell lines containing either the GFP reporter or HA-tagged protein VII under a doxycycline-inducible promoter were generated as previously reported^32,66^. After plasmid transfection and puromycin selection, HEK293T HILO cell lines were grown in DMEM (Fisher #11965118) with tetracycline-free 10% FBS (Gemini Bio-Products #100-800; Lot #A69G00J), 1% penicillin-streptomycin (Fisher #15-140-122) and 1 ug/mL puromycin (Fisher #MIR5940) to maintain selection.

### Exogenous Protein Expression in Human Cells

RPE-1 cells were plated at 20% confluence in 6-well plates and the following day, cells were transduced with recombinant adenovirus with the E1 region replaced with either GFP-tagged protein VII (MOI = 50) or a GFP control (MOI = 50) under a CMV promoter as previously described^90^. Control cells were treated with 10 μM RO-3306 (Sigma-Aldrich #SML0569-5MG). For all treatment groups, samples were collected every 12 hours post-treatment for 48 hours total as described below. The HEK293T doxycycline-inducible cell lines were removed from puromycin selection and plated at a density of 2.6 x 10^4^ cells/cm^2^ in 60 mm plates with 0.2 μg/mL doxycycline to induce protein expression. All cell lines and treatment groups were imaged, and samples were collected every 24 hours post-treatment for 96 hours. Cells were imaged using Leica Microsystems DM IL LED Fluorescent microscope with Leica Application Suite V4.12 software. Images were processed using FIJI, Adobe Photoshop, and compiled with Adobe Illustrator.

### Human Cell Proliferation Assay

RPE-1 cells were harvested every 12 hours after transduction for 48 hours in addition to a pre-transduction sample. HEK293T-based cells were harvested every 24 hours after plating and induction up to 96 hours. For each collection, both cell types were lifted by trypsinization. Cells were pelleted by centrifugation (1200 rpm for 2 minutes) and then suspended in 1 mL DPBS + 1 mM EDTA to prevent clumping. 20 uL cell suspension was diluted 1:1 with 0.4% Trypan Blue solution (Bio-Rad #1450021) and cells were counted by hemocytometer (RPE-1 cells) or using a BioRad TC20 Automated Cell Counter (HEK293T-based cells lines). The total number of cells in each well and the number of trypan negative cells was reported.

### Human Cell Flow Cytometry

Cells were lifted with trypsin, pelleted by centrifugation (1000 rpm for 5 minutes at 4°C), and washed in 5 mL FACS Buffer (DPBS + 0.1% BSA). Cells were pelleted and resuspended in 1 mL 0.5% PFA diluted in DPBS and incubated for 5 minutes at room temperature to fix the cells. Cells were washed twice in FACS Buffer and kept on ice or at 4°C for all remaining processing. Cells were then suspended in 70% ethanol and stored at −20°C overnight. Next, the cells were washed three times in DPBS + 0.1% Triton-X, then resuspended in 500 uL DPBS + 0.2 mg/mL RNAse A (Sigma # 10109142001) and 20 ug/mL propidium iodide (Fisher # AC440300250), then incubated in the dark at 4°C for at least 3 hours. Cells were run on a BD Biosciences FACSCanto II Cell Analyzer in the Fred Hutchinson Cancer Research Center Flow Cytometry core facility. Since GFP was present in >90% of the cells expected to express GFP, cells were not gated based on GFP. Instead, single cells were gated using propidium iodide area and height measurements. Histograms indicating DNA content were generated using FlowJo v10 software.

### Human Cell Western Blotting

Western blot cell samples were counted and then pelleted and resuspended in 1x NuPAGE LDS sample buffer (Fisher #NP0007) + 5% 2-betamercaptoethanol. The mixture was heated for 20 minutes at 95°C and stored at −20°C. 10 uL of each sample was run on a 15% SDS PAGE gel using standard methods as described above with the following primary antibodies: α-vinculin (1:20,000 Sigma #V9131), α-GFP (1:2500, Abcam #290), α-H3 (1:20,000, Abcam #1791, α-H3S10ph (1:1000, EMD Millipore #06-570), α-HA (1:500, BioLegend/Covance #MMS-101R) or α-E1A (1:250, BD Biosciences #554155); and secondary antibodies: α-mouse (Jackson ImmunoResearch #115-035-003) or α-rabbit (Jackson ImmunoResearch #111-035-045). Blots were developed with Clarity Western ECL (Bio-Rad #1705061) and images were processed as described above.

### Supplemental Figures and Table

**Supplemental Figure 1.**
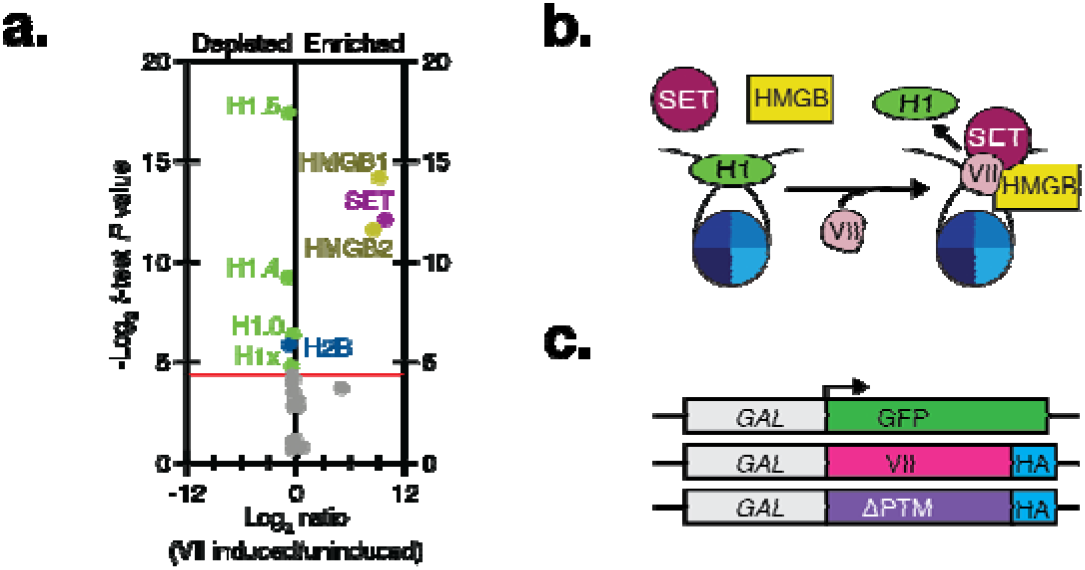
Binding of the histone-like adenovirus protein VII to chromatin causes linker histone depletion. **a.** Volcano plot of mass spectrometry analysis of salt-fractionated chromatin from cells with and without protein VII expression from our previously published study^32^. The data points correspond to protein SET (purple), HMGB proteins (gold), linker H1 (green), and all detected core canonical histones (blue, gray). The y-axis represents the −log_2_ statistical *p-*value and the x-axis represents the log_2_ protein fold-change between uninduced or protein VII-expressing cells (homoscedastic two-tailed t-test, *p* < 0.05) indicating whether a protein was enriched or depleted from chromatin. The points shown in gray fall below the cutoff for statistical significance which is indicated by the red line. See Supplemental Table 1 for all protein identifiers and statistical values. **b.** Model for protein VII disruption of chromatin in human cells. Proteomics of protein VII-bound chromatin suggests its interaction with HMGB1 and SET facilitate protein VII invasion of chromatin and displacement of linker histone H1. *In vitro,* protein VII binds the nucleosome at a position similar to linker histones suggesting that H1 and VII compete for binding the nucleosome. **c.** Schematics of budding yeast expression construc**ts.** The galactose promoter is truncated to drive transcription only in the direction indicated by the arrow. The plasmids code for either GFP, mature protein VII from WT adenovirus type 5, or protein VII with alanine substitutions at 5 sites of post-translational modification (ΔPTM). VII and VII-ΔPTM genes are codon optimized for expression in *S. cerevisiae* and contain a 3xHA epitope tag at the C-terminus. Each of these promoter and gene combinations is integrated into the same *URA3-*selectable plasmid backbone with a centromere to ensure high fidelity of transmission.

**Supplemental Figure 2.**
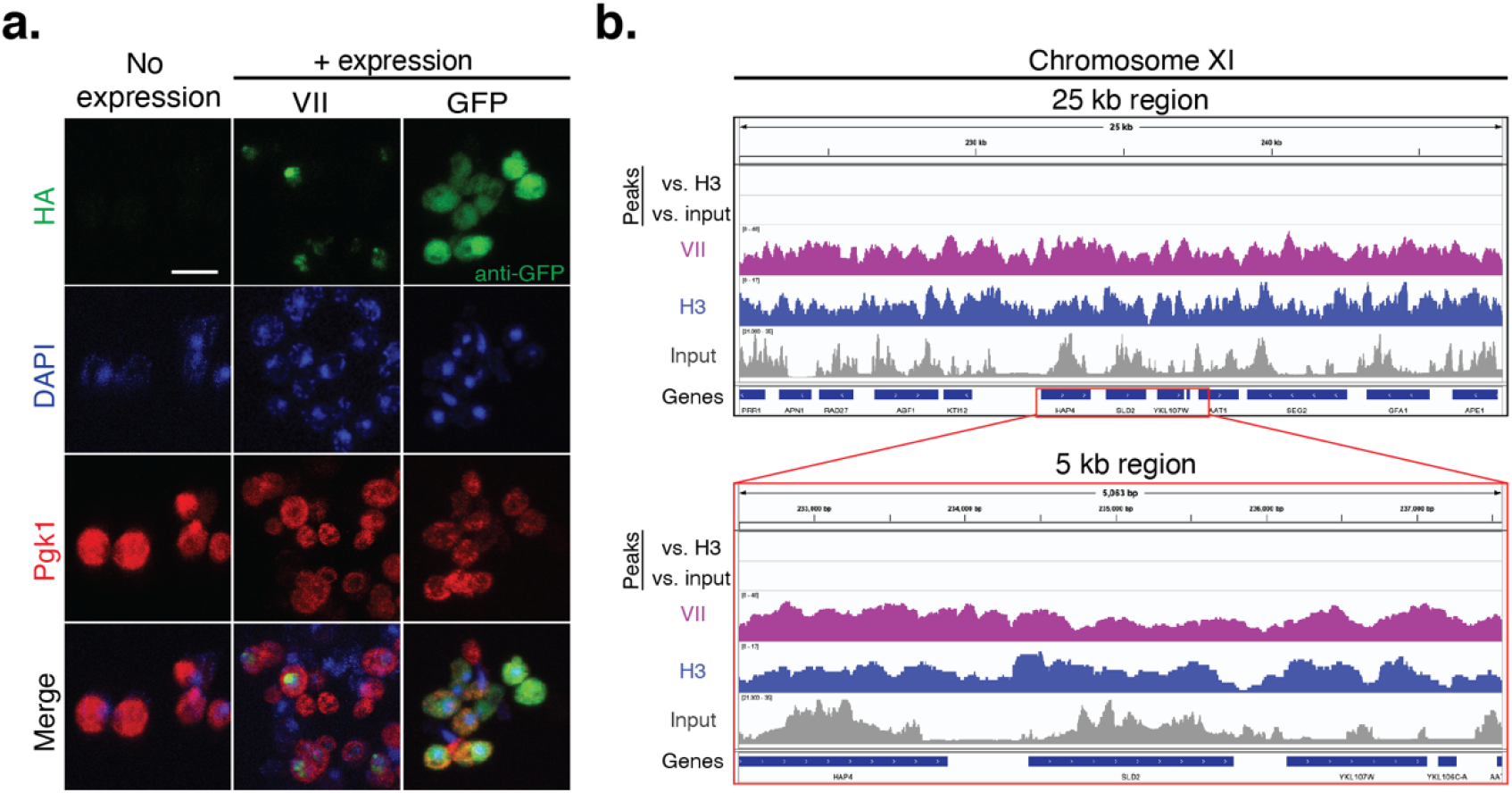
Protein VII localizes to the nucleus and binds chromatin in yeast. **a.** Immunofluorescence microscopy of WT FY602 cells with and without protein expression. Pgk1 indicates the cytoplasm and DAPI stains the nucleus. Scale bar is 5 μm. **b.** Native ChIP-seq of WT FY602 cells expressing protein VII. “Narrow peaks” called by MACS2 peak caller are shown in the top two tracks along with IGV tracks for input, histone H3, and protein VII ChIP-seq reads at one 25 kb locus on Chromosome XI. Red box indicates a smaller 5 kb region shown in greater detail. All tracks were generated with MACS2 peak-caller using uniquely mapped reads. Reads are normalized to counts per million. Tracks are automatically scaled for each chromosome. Tracks from one experimental replicate are shown. ChIP-seq was performed in duplicate.

**Supplemental Figure 3.**
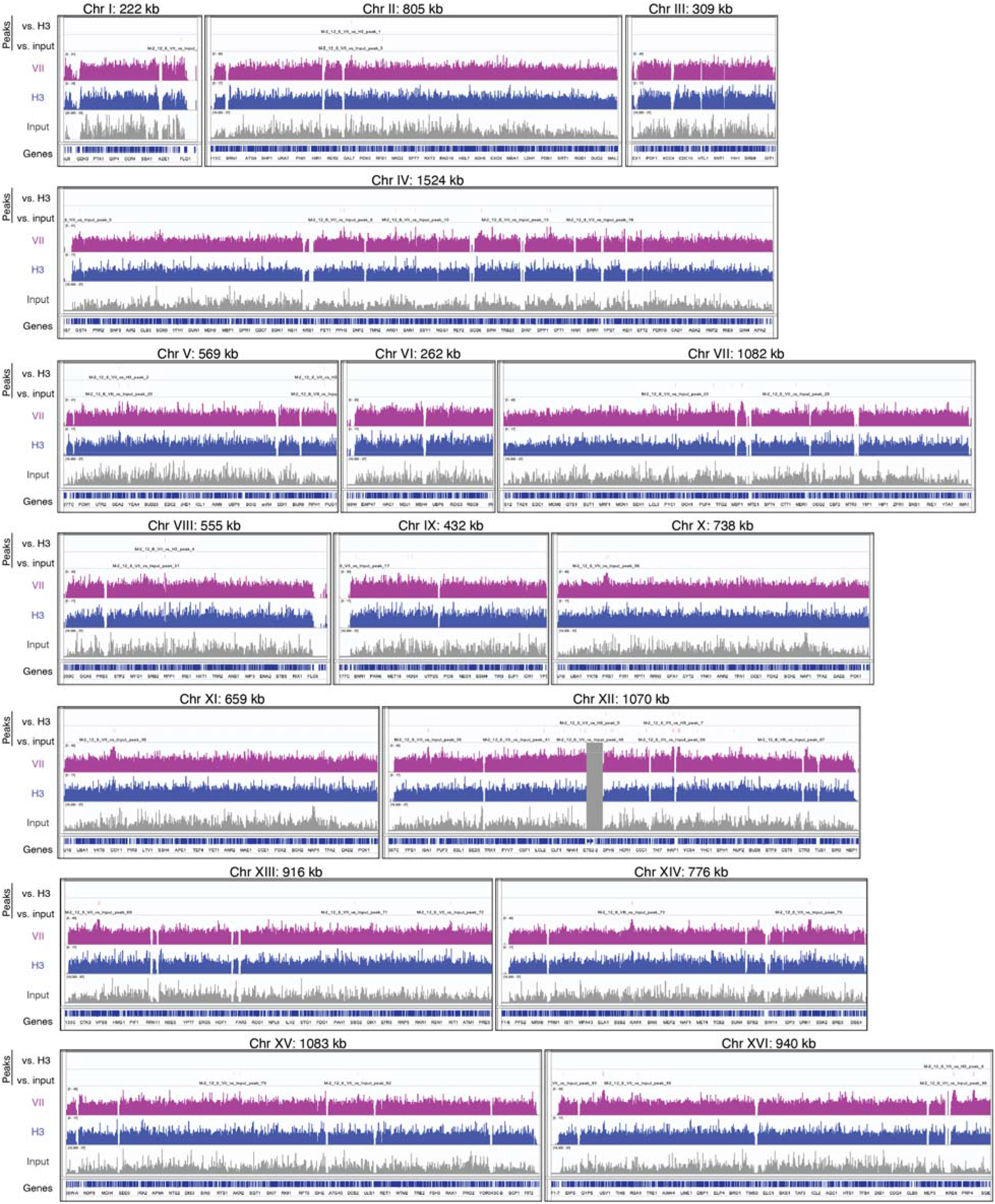
Protein VII binds throughout yeast chromatin. “Narrow peaks” called by MACS2 peak caller are shown along with IGV tracks for input, histone H3, and protein VII ChIP-seq for all yeast chromosomes from WT FY602 as described in Supplemental Figure 2b. Due to poor mapping, the reads at the rDNA locus (chromosome XII; ~450 – 480 kb) are masked.

**Supplemental Figure 4.**
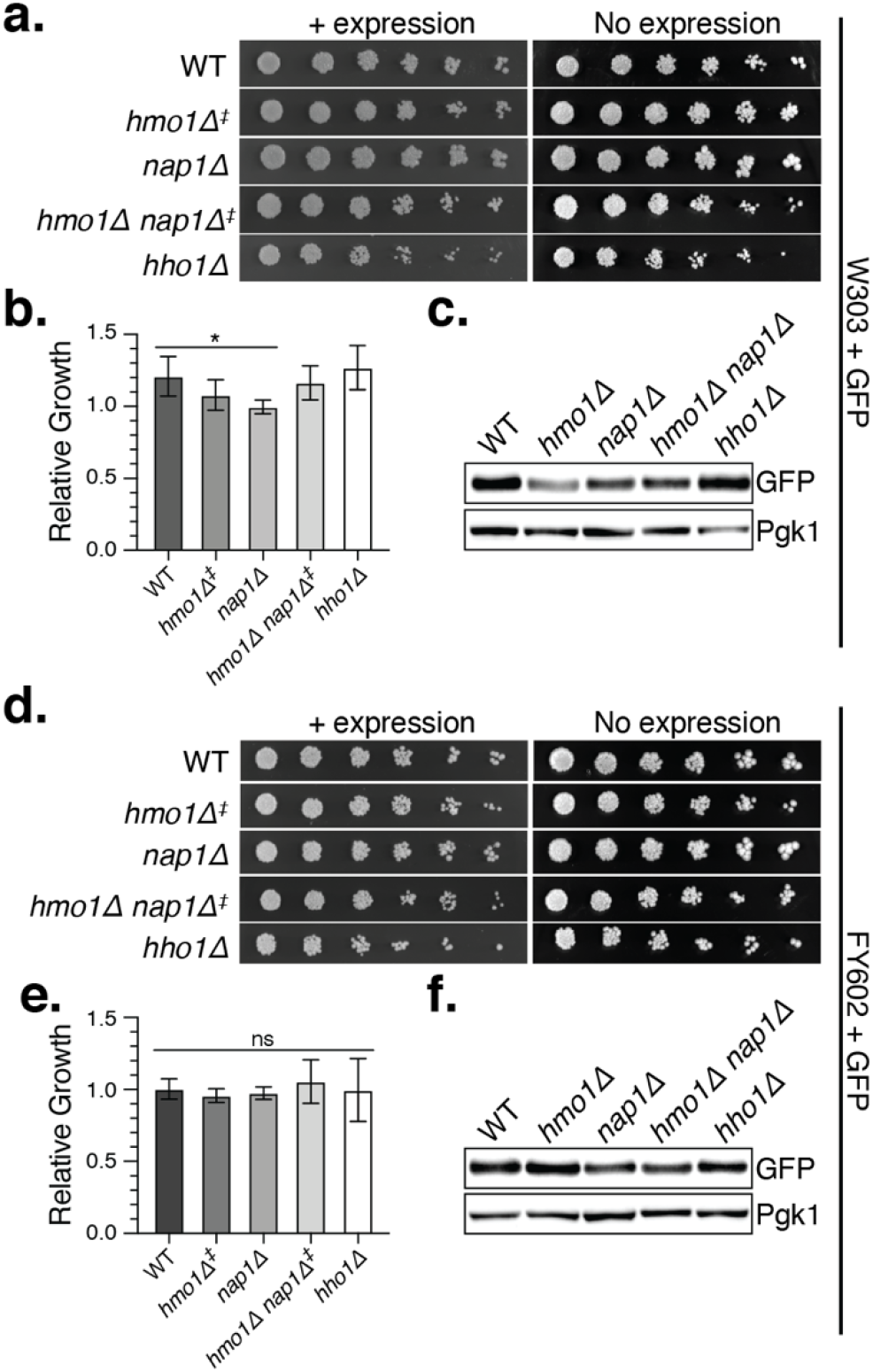
Loss of Hmo1 or Nap1 has no significant impact on growth when GFP is expressed. **a.** Representative serial dilution assay for growth defects of induced and repressed W303 cells with GFP expression plasmid in the indicated gene deletion strains as in Figure 2b. **b.** Quantification of serial dilution growth assay in W303 gene deletion strains during GFP expression. Relative growth was quantified as in Figure 2c for at least three biological replicates. No statistical significance was found when WT growth was compared to each deletion strain except for the *nap1Δ* strain, although the *p*-value barely exceeded the cut-off for significance (One-way ANOVA with multiple comparisons vs. WT; *p*=0.0443). *P*-values for remaining comparisons to WT: *hmo1Δ* = 0.3213, *nap1Δ* = 0.0443, *hmo1Δnap1Δ* = 0.9583, *hho1Δ* = 0.8666. Error bars represent SD. **c.** Western blot analysis of induced W303 gene deletion strains with GFP expression plasmid and Pgk1 as a loading control. **d.** Representative serial dilution assay for growth defects in induced and repressed FY602 cells with GFP expression plasmid in the indicated gene deletion strains as described in Figure 2b. **e.** Quantification of serial dilution growth assay in FY602-based deletion strains when GFP is expressed as described in Figure 2c. No statistical significance was determined among the deletion strains when compared to WT growth using one-way ANOVA with multiple comparisons. *P*-values for comparisons vs. WT: *hmo1Δ* = 0.8159, *nap1Δ* = 0.9723, *hmo1Δnap1Δ* = 0.8202, *hho1Δ* = 0.9998. Error bars depict SD. At least four biological replicates were quantified for each strain. **f.** Western blot analysis of induced WT FY602 cells and related deletion strains with GFP expression plasmid and Pgk1 shown as the loading control.

**Supplemental Figure 5.**
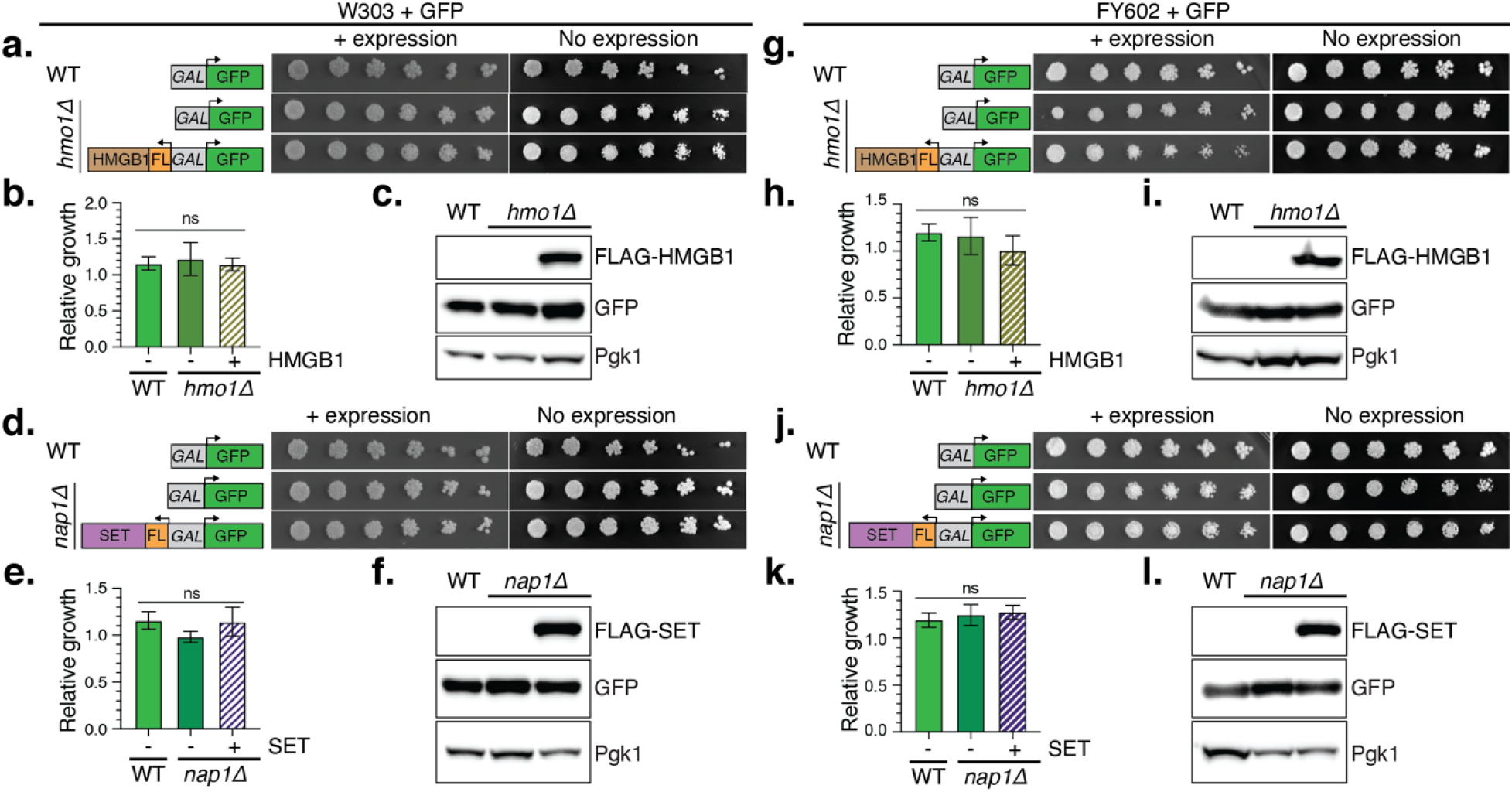
Replacing Hmo1 or Nap1 with homologous human factors has no significant impact on growth when GFP is expressed. **a.** Serial dilution spot assay for GFP control in WT W303 and W303 *hmo1Δ* strain with and without co-expression of human HMGB1. Expression constructs illustrated to the right show the bidirectional galactose-inducible promoter that drives GFP and human HMGB1 transcription. Cells without protein expression are shown after 3 days growth at 30°C while cells with protein expression are shown after 5 days growth at 30°C to compensate for slowed growth on the non-preferred energy source, galactose. **b.** Quantification of serial dilution growth assay of strains shown in (a). Growth was quantified after 5 days growth with protein expression and compared to growth after 3 days growth without protein expression. A one-way ANOVA with multiple comparisons showed no significant difference in growth among the three different strains and co-expression constructs (WT + GFP vs. *hmo1Δ* + GFP *p*=0.8426; WT + GFP vs. *hmo1Δ* + GFP + HMGB1 *p*=0.9922, *hmo1Δ* + GFP vs. *hmo1Δ* + GFP + HMGB1 *p*=0.8205). Error bars represent SD for all growth quantification in this figure. N= 3. **c.** Western blot analysis of strains shown in (a). FLAG-HMGB1, GFP, and Pgk1 loading control are shown. **d.** Serial dilution growth assay for GFP control in WT W303 and W303 *nap1Δ* strain with and without co-expression of human SET. Bidirectional galactose promoter and target genes are illustrated at right. Images depicting growth were selected as described in (a). **e.** Quantification of serial dilution growth assay in WT W303 and W303 *nap1Δ* when GFP is expressed alone or co-expressed with human SET as described in (b). A one-way ANOVA with multiple comparisons failed to detect statistically significance differences in growth among these strains (WT + GFP vs. *nap1Δ* + GFP *p*=0.1181; *nap1Δ* + GFP + SET *p*=0.9832, WT + GFP vs. *nap1Δ* + GFP + SET *p*=0.1536). N= 3. **f.** Western blot analysis from strains shown in (d). GFP and FLAG-SET blots are shown with Pgk1 loading control. **g.** Serial dilution growth assay for GFP expression in WT FY602 and FY602 *hmo1Δ* strain with and without co-expression of human HMGB1 as described in (a). **h.** Quantification of serial dilution growth assay in WT W303 and W303 *hmo1Δ* when GFP is expressed alone or co-expressed with human HMGB1 as described in (b). There was no significant difference in growth among the three strains (one-way ANOVA with multiple comparisons; WT + GFP vs. *hmo1Δ* + GFP *p*=0.9517; WT + GFP vs. *hmo1Δ* + GFP + HMGB1 *p*=0.3470, *hmo1Δ* + GFP vs. *hmo1Δ* + GFP + HMGB1 *p*=0.4853). N= 3. **i.** Western blot analysis of WT FY602 with GFP, *hmo1Δ* with GFP, and *hmo1Δ* with GFP and HMGB1 co-expression. FLAG-HMGB1, GFP, and Pgk1 loading control blots are shown. **j.** Serial dilution growth assay for GFP in WT FY602 and FY602 *nap1Δ* strain with and without co-expression of human SET as described in (d). **k.** Quantification of serial dilution growth assay in WT W303 and W303 *nap1Δ* when GFP is expressed alone or coexpressed with human SET as in (b). There was no significant difference in growth among the three strains (one-way ANOVA with multiple comparisons; WT + GFP vs. *nap1Δ* + GFP *p*=0.7047; WT + GFP vs. *nap1Δ* + GFP + SET *p*=0.4743, *nap1Δ* + GFP vs. *nap1Δ* + GFP + SET *p*=0.9222). N= 3. **l.** Western blot analysis of strains shown in (j). FLAG-SET, GFP, and Pgk1 loading control blots are shown.

**Supplemental Figure 6.**
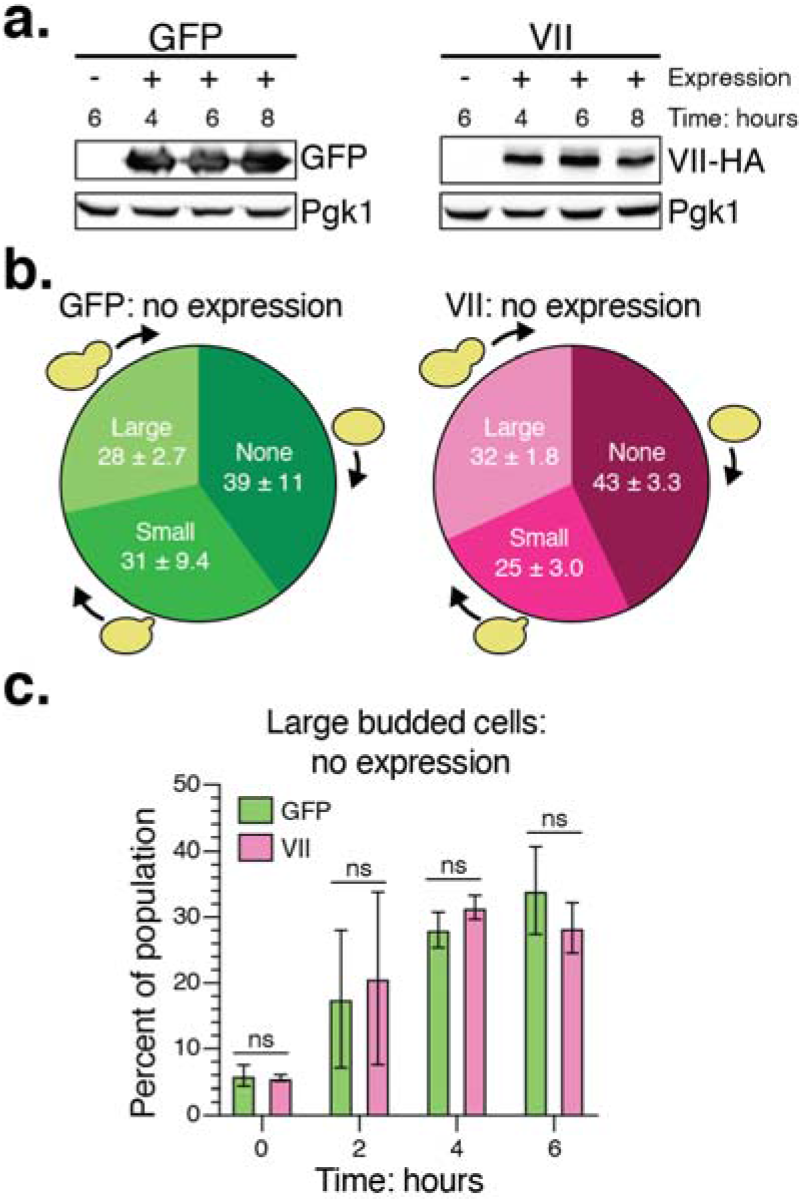
The accumulation of large-budded cells requires protein VII expression. **a.** Western blot analysis of timed samples collected during budding analysis of WT W303. Time indicates total time in hours that the cells grew in the no expression (dextrose) or + expression (galactose) condition. GFP, VII-HA, and Pgk1 loading control blots are shown. **b.** Budding analysis of strains shown in Figure 4 without induction of protein expression and after 4 hours asynchronous growth in dextrose media. The percentage of the population with each bud type is shown along with SD. Multiple unpaired t-tests failed to identify a statistically significant difference in the proportion of cells in each budding class between the two strains (*p*=0.6284 for unbudded cells, *p*=0.3329 for small-budded cells, and *p*=0.1445 for large-budded cells). N=3 in WT W303. **c.** The percentage of large-budded cells in the population during asynchronous growth without induction of GFP (green) or protein VII (pink) expression. At the beginning of the experiment, the cells in stationary phase (T0) were moved to dextrose media to repress protein expression and were grown to mid-log phase (T6). The s-axis indicates the time since the addition of dextrose and the start of the growth experiment. Unpaired *t-*test *P*-values for each timed sample in order as shown on the graph from T0 to T6 are: 0.7271, 0.7601, 0.1445, and 0.2701. N=3 in WT W303.

**Supplemental Figure 7.**
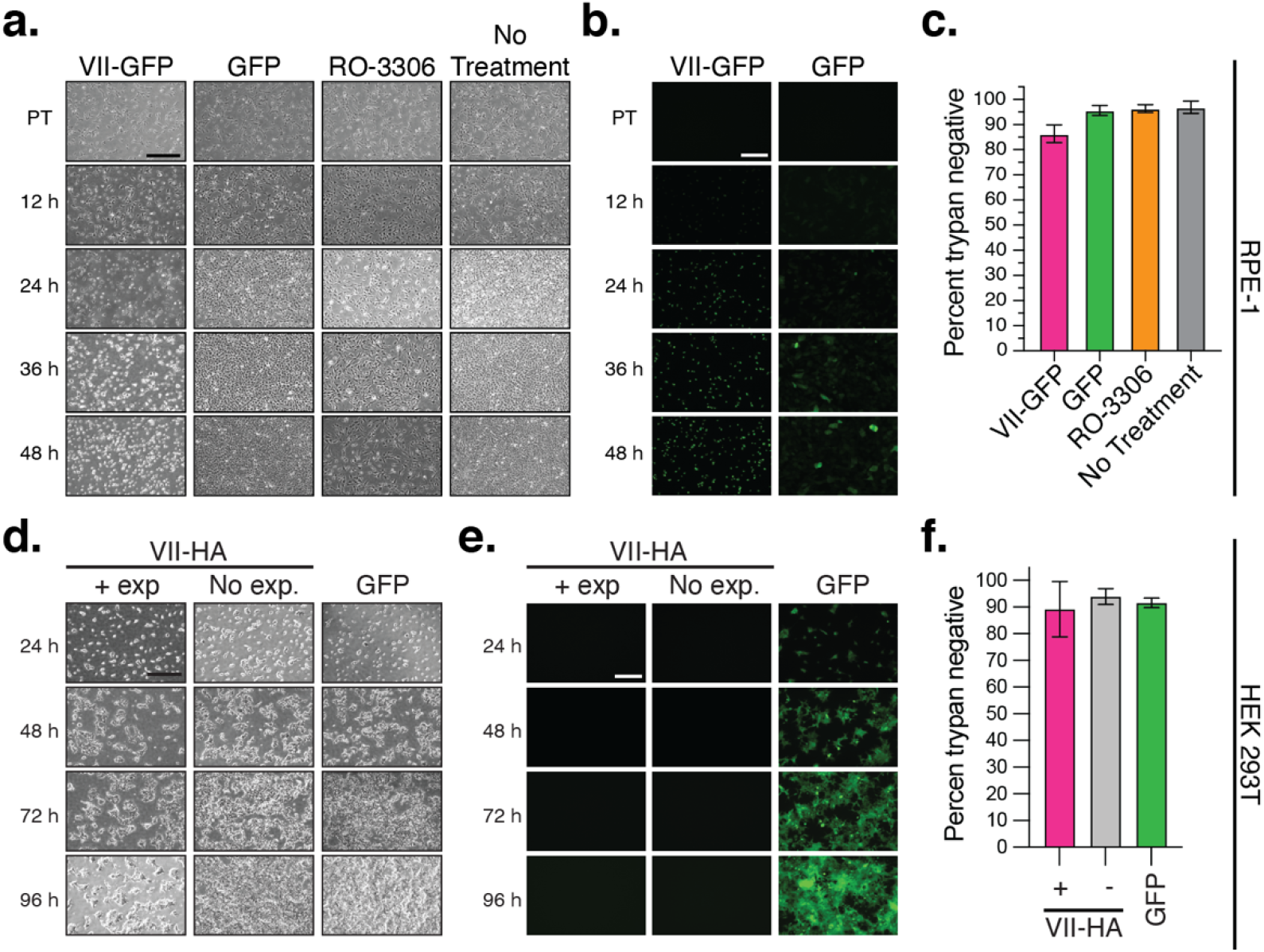
Non-proliferative cells that express protein VII are metabolically active. **a.** Brightfield microscopy images of diploid RPE-1 cells show the loss of proliferation in cells transduced to express protein VII-GFP, or treated with RO-3306, compared to the no treatment and GFP controls. 5x magnification. Scale bar is 500 μm. **b**. Fluorescent microscopy images of RPE-1 cells that express either protein VII-GFP or GFP. 10x magnification. Scale bar is 200 μm. **c**. Percentage of RPE-1 cells that are negative for trypan blue exclusion stain 48 hours post-treatment. Errors bars are SD. N = 4. **d**. Brightfield microscopy images of HEK 293T cells show the loss of proliferation in cells expressing protein VII-HA but not the untreated control or cells expressing GFP. 5x magnification. Scale bar is 500 μm. **e**. Green fluorescent microscopy images of HEK 293T cell lines with and without doxycycline induction of protein expression. 10x magnification. Scale bar is 200 μm. **f**. Percentage of HEK 293T cells negative for trypan blue exclusion stain 96 hours post-treatment. indicates doxycycline treatment status of VII-HA cells. The GFP cells are induced with doxycycline. Errors bars are SD. N = 3.

**Supplemental Table 1.** A subset of protein and gene identifiers as well as statistical and enrichment scores for proteomics of salt-fractionated chromatin from cells with and without protein VII expression as previously published^32^. Here, the *t*-test *p*-values and enrichment scores corresponding to protein SET, HMGB proteins, linker H1 types, and the core canonical histones are shown. The values shown in gray-filled boxes were not statistically significant (*p*<0.005).

| Protein Name | Gene Names | *t*-test *P*-value | Log2 fold change: VII induced/uninduced |
| --- | --- | --- | --- |
| Histone H1.5 | HIST1H1B | 5.69851E-06 | -0.757738748 |
| High mobility group protein B1; Putative high mobility group protein B1-like 1 | HMGB1; HMGB1P1 | 5.38059E-05 | 9.17634819 |
| Protein SET | SET | 0.000229685 | 9.848560151 |
| High mobility group protein B2 | HMGB2 | 0.000322399 | 8.559909323 |
| Histone H1.4 | HIST1H1E | 0.001652512 | -0.903455652 |
| Histone H1.0; Histone H1.0, N-terminally processed | H1F0 | 0.012110662 | -0.359847618 |
| Histone H2B type 3-B | HIST3H2BB | 0.017135066 | -0.68225997 |
| Histone H1x | H1FX | 0.035053385 | -0.495869309 |
| Histone H2A type 1-C; Histone H2A type 3; Histone H2A; Histone H2A type 1-B/E | HIST1H2AC; HIST3H2A; HIST1H2AB | 0.05001239 | -0.398885931 |
| Histone H1.2 | HIST1H1C | 0.060260505 | -0.410806769 |
| Histone H3; Histone H3.3 | H3F3B; H3F3A | 0.076305074 | 5.001910361 |
| Histone H4 | HIST1H4A; HIST1H4L; HIST1H4H | 0.086418311 | -0.304855891 |
| Histone H2A.V; Histone H2A.Z; Histone H2A | H2AFV; H2AFZ | 0.119555025 | 0.216370185 |
| Histone H2A type 2-C; Histone H2A type 2-A | HIST2H2AC; HIST2H2AA3 | 0.140693297 | -0.2801282 |
| Histone H2B type 1-M; Histone H2B type 1-N; Histone H2B type 1-H; Histone H2B type 2-F; Histone H2B type 1-C/E/F/G/I; Histone H2B type 1-D; Histone H2B; Histone H2B type 1-K; Histone H2B type F-S | HIST1H2BM; HIST1H2BN; HIST1H2BH; HIST2H2BF; HIST1H2BC; HIST1H2BD; HIST1H2BI; HIST1H2BK; H2BFS | 0.144611424 | 0.26332697 |
| Histone H2A.x; Histone H2A type 1-A | H2AFX; HIST1H2AA | 0.146404666 | -0.185106615 |
| Histone H2B type 2-E; Histone H2B type 1-B; Histone H2B type 1-O; Histone H2B type 1-J | HIST2H2BE; HIST1H2BB; HIST1H2BO; HIST1H2BJ | 0.456113639 | -0.103554106 |
| Histone H2B type 1-L; Histone H2B type 1-A | HIST1H2BL; HIST1H2BA | 0.479413333 | -0.409269021 |
| Histone H3.2 | HIST2H3A | 0.574677306 | -0.226071755 |
| Histone H3.1; Histone H3.1t | HIST1H3A; HIST3H3 | 0.584639377 | 0.694148161 |
| Histone H1.3 | HIST1H1D | 0.633380729 | -0.344162535 |

